# Wild *Luteibacter* populations exhibit climate-associated divergence and distinct genomic strategies for osmotic stress tolerance

**DOI:** 10.64898/2026.09.27.754828

**Authors:** Nichole Ginnan, Carmen Rodriguez, Brian J. Sanderson, Felicity Tso, Natalie E. Ford, Manuel Kleiner, Maggie R. Wagner

## Abstract

Soil bacteria play a crucial role in supporting ecosystem health, yet little is known about how wild populations adapt to abiotic stressors, like limited water availability. We used 186 *Luteibacter* isolates from maize roots grown in never-irrigated prairie soils collected from a steep precipitation gradient in Kansas, US to test whether climate history is associated with osmotic stress tolerance and genomic differentiation. *In vitro*, strains from semi-arid sites were more tolerant to osmotic-stress treatment than those from humid sites. Comparative genomics of a subset of 96 isolates identified four strongly biogeographically structured lineages and extensive accessory-genome diversity. Semi-arid climate-associated strains were enriched for genes involved in osmotic homeostasis, oxidative stress protection, motility, and type IV secretion. Humid climate-associated strains were enriched in environmental sensing, metabolic versatility, biofilm formation, iron acquisition, type VI secretion, and resource competition. GWAS identified 22 candidate genes associated with osmotic-stress tolerance, including those related to type IV pili, biofilm production, DNA repair, and phage. No candidates were shared between semi-arid and humid strains, suggesting distinct genetic mechanisms of osmotic tolerance. Predicted prophage composition was also geographically structured, although total prophage gene content did not predict osmotic-stress tolerance. Together, these results show that adaptation to long-term differences in water availability likely involves coordinated shifts in stress physiology, genome content, and ecological interactions in wild bacterial populations, providing insight into predicting microbial responses to changing climates.

**IMPORTANCE:** Soil bacteria support plant growth, nutrient cycling, and ecosystem health, but we know surprisingly little about how natural bacterial populations adapt to different climates. By studying hundreds of *Luteibacter* strains collected across a natural precipitation gradient, we show that bacteria from drier environments are better able to tolerate water-related stress, but this resilience comes with trade-offs in growth. We also found that populations from dry and wet climates use different genetic and ecological strategies to cope with stress, including differences in metabolism, cell-surface traits, and interactions with bacterial viruses. These findings demonstrate that bacterial adaptation to climate involves more than a single stress-response mechanism. Understanding this natural diversity can improve predictions of how soil microbiomes will respond to climate change and help identify or engineer resilient microbes for agricultural and environmental applications.

## INTRODUCTION

Soils harbor more than 50% of global biodiversity, and the majority of those lifeforms are microbial (1). These complex soil microbial communities perform essential functions, like nutrient cycling (2) and carbon sequestration (3), and are major determinants of soil, plant, and broader ecosystem health (4). Climate change poses an unprecedented threat to the persistence of life across scales (5–7). Bacteria, many of which can adapt more rapidly than more complex organisms due to their relatively short generation times, are increasingly studied for their potential to mitigate the negative impacts of climate stressors, particularly drought (8, 9). Droughts are expected to increase in frequency and intensity, and understanding how soil bacterial populations, within complex natural soil communities, adapt to osmotic stress is important for prediction and alleviation of future drought effects on soil microbiomes.

Soil bacteria live in highly dynamic environments and need to withstand persistent environmental stresses and rapid fluctuations, particularly shifts in osmolarity and osmotic pressure related to weather events and seasonal changes (10). To survive under drought or low water availability, bacteria firstly rely on accumulating compatible solutes that regulate internal osmotic pressure, such as proline and glycine betaine (11, 12). Secondly, bacteria activate stress-response genetic programs to launch a physiological response (11). As gene expression shifts toward the production of stress-responsive proteins, bacterial growth slows, which is critical for persistence under environmental stress. Therefore, measuring and comparing the reduction in growth is an informative measure of sensitivity to a stressor (11). Differences in cell envelope structure (12, 13) and biofilm formation (14, 15) are commonly associated with variation in osmotic-stress tolerance. Notably, stress tolerance is a complex, multigenic trait (16), and bacteria often exhibit cross-adaptation, where resistance to one stressor also confers tolerance to others through shared mechanisms (17, 18). This means that dry climate adaptations could also affect tolerance to other biotic and abiotic stressors (19). However, because cellular resources are finite, investment in stress adaptation can reduce growth efficiency, potentially resulting in a trade-off between survival and proliferation (20). Bacterial genetic adaptations to osmotic stress have primarily been studied under laboratory conditions or in model organisms (12, 13), leaving a significant gap in our understanding of wild microbial adaptive responses to abiotic stress (21). More specifically, it is unclear if native soil bacterial populations adapt to abiotic stress in the same way as lab-grown populations, and which genetic mechanisms underlie this adaptation.

Here, we investigate osmotic stress tolerance of a soil bacterial genus from never-irrigated prairies that can also readily colonize plant roots, *Luteibacter* spp. The genus *Luteibacter*, of the *Rhodanobacteraceae* family within Gammaproteobacteria (22), can be high in relative abundance in both bulk soil and plant rhizosphere communities (23) and becomes enriched in the wheat root endosphere under drought conditions (24). *Luteibacter* can stimulate barley root growth (25), and was a constituent of a 15-member synthetic bacterial community that enhanced *Brachypodium distachyon* plant growth under drought-rewetting and salinity stress (26). Although *Luteibacter* remains an under-researched genus, these findings indicate that *Luteibacter* spp. thrive in diverse environments, including a variety of host plants, and exhibit resilience to water stress with the potential to enhance drought tolerance in plants.

We sampled strains of *Luteibacter* across a well-characterized precipitation gradient in Kansas, USA, where soils in the east have a history of exposure to high precipitation (humid climate) and western soils experience low precipitation (semi-arid climate) (27, 28). This natural gradient provides a powerful framework for testing how abiotic stressors shape soil bacterial genomes and traits. Previous work using *16S rRNA* gene and shotgun metagenomic sequencing indicates that bacterial community structure and function vary across precipitation gradients (28, 29). However, metagenomic sequencing has limited ability to provide population-level resolution and identify genetic drivers of bacterial local adaptations in complex systems, like soil (30). Additionally, genome-wide association studies (GWAS) are useful tools for phenotype-to-genotype mapping, and establishing a microbial phenotype is significantly easier if the strains are in pure culture. Moreover, the high variability of bacterial genomes (HGT, insertions, deletions, etc.) and potentially high dispersal rates that might homogenize soil bacterial populations have limited our understanding of environmental bacterial genetic adaptations (31–34).

We used *Luteibacter* as a model for understanding if wild soil-dwelling bacteria associated with different environmental conditions differ in osmotic stress tolerance, and if so, identify genetic factors potentially driving differential trait expression. We hypothesized that 1. *Luteibacter* strains from semi-arid soils would be more tolerant to osmotic stress than strains from humid soils, 2. Climate history would be associated with genome-wide patterns of genetic differentiation, and 3. candidate genetic drivers of bacterial osmotic stress tolerance could be identified using a microbial GWAS approach. To test this, we screened strains using a quantitative growth assay under increasing salt (NaCl) concentrations, which induces both osmotic and ionic stress (35), and polyethylene-glycol (PEG), which induces only osmotic stress (36). Next, we assembled a pangenome of a subset of our strains and identified genomic features associated with osmotic stress tolerance. We then used a microbial GWAS approach tailored for structured bacterial populations (37) to identify genetic variants associated with stress tolerant phenotypes. Interestingly, we found fitness trade-offs between growth and osmoadaptation in our bacterial population.

## MATERIALS & METHODS

### Bacterial isolates collection

Pure *Luteibacter* isolates were selected from a larger maize root-associated bacterial culture collection described previously (Supplemental Methods)(38). Specifically, we used *Luteibacter* strains that originated from prairie soils from four Kansas, USA sites spanning a precipitation gradient: The Land Institute (TLI), Konza Prairie Biological Station (KNZ), Hays Prairie (HAY), and Smoky Valley Ranch (SVR). Microbial slurries from these soils were inoculated onto sterile maize seeds, and bacteria were subsequently isolated from one-month-old maize roots (Supplemental Methods)(38). Isolates were initially identified by sequencing of the full length *16S rRNA* gene. From this effort we obtained 549 *Luteibacter* spp. isolates, which we used as the starting population for this study.

### Kansas precipitation gradient map

Average temperature data from Kansas counties for years 2010-2020 was downloaded from NOAA’s National Centers for Environmental Information website (39) (Supplemental Table S1). To visualize spatial patterns in precipitation across Kansas, mean annual precipitation (2010–2020) for each county was calculated from raw climate data using dplyr (v1.1.4) package in R (40). County-level shapefiles for Kansas were obtained using tigris (v2.2.1) (41) and were joined to the precipitation values. Geographic coordinates for each soil sampling site were converted to spatial points with sf (v1.0-21) (42, 43), and mapped onto the county shapefile. The map was generated using ggplot2, with mean precipitation displayed as color gradient and sampling sites overlaid and labeled.

### Preliminary qualitative screening

First, to identify a subset of strains that varied in stress tolerance for follow-up quantitative assays and whole genome sequencing, we conducted a preliminary *in vitro* salt assay on R2A medium. Concentrations of 0%, 0.3%, 1.8%, and 3% NaCl were mixed with R2A agar, autoclaved, then poured onto rectangular petri plates. *Luteibacter* cultures were stamped onto the solid medium using a flame sterilized microplate replicator dipped into the 96 well plate glycerol stocks. These plates were incubated at 28°C for 48 hours and then visually evaluated for growth at the varying NaCl concentrations (Supplemental Fig. 1). The most tolerant (growth on 1.8% and 3% NaCl) and the most sensitive (no growth on 0.3%) strains within each site were selected for sequencing (96 strains) and additional phenotyping (186 strains). In addition to this qualitative phenotyping, the *16S rRNA* gene tree details were incorporated into the stain selection to maximize phenotypic and genetic diversity (Supplemental Fig. 2).

### 16S rRNA gene tree

The full-length *16S rRNA* gene sequences of the 549 *Luteibacter* isolates were aligned using MUSCLE v3.8 (44). The multiple sequence alignment was trimmed using ClipKIT (mode=smart-gap) (45). A maximum likelihood tree was built using MEGA 12 (model=Tamura-Nei (46)) (47). The tree and tree metadata were visualized using iTOL (48) (Supplementary Fig. S2).

### *Q*uantitative osmotic stress phenotyping of select *Luteibacter* strains

Of the 549 *Luteibacter* isolates, 186 strains were selected for screening based on their *16S rRNA* gene diversity within each original soil and their phenotypes in the initial qualitative assay. Each selected bacterial strain was grown for 48 hours at 29°C on R2A agar medium. Single isolated colonies of each strain were inoculated into individual tubes of R2A liquid medium. The inoculants were incubated at 29°C, shaking at 250 rpm for 24 hours. In a 96 well plate, 500 µl of liquid culture and 500 µl of 50% glycerol were added into each well in randomized order with three replicates of each strain, as well as three blank uninoculated wells. Overall, there were six 96-well plates containing 31 different isolates each, including three replicates of each strain. Plates were sealed and stored at −80°C.

First, the selected 186 *Luteibacter* strains were grown in standard liquid R2A medium to measure cell growth in the absence of osmotic stress. Using 96-well black clear-bottomed assay plates (Corning Incorporated, Corning, NY USA; #3603), 200 µl of liquid R2A media was added into each well. Approximately 1 µl of bacterial culture (including glycerol) was transferred from the 96-well plates into the assay plates using a flame-sterilized microplate replicator. Plates were covered with Aeraseal films (Excel Scientific, St. Paul, MN, USA; #BS-25) and lids, then incubated at 29℃ shaking at 250 rpm for 24 hours. Then, the liquid culture in each well was mixed by aspirating and dispensing using a pipette. Next, 100 µl was removed from each well and discarded, leaving 100 µl of culture in each well. Under dark conditions, 20 µl of Promega CellTiter-Blue reagent (Fisher Scientific, Waltham, MA, USA; #PR-G8081) was added into each well and covered with tin foil to avoid light exposure. The cultures were placed back into the incubator at 29℃ and shaken at 250 rpm for 4 hours to develop. A CLARIOstar Plus microplate reader (BMG LABTECH, Cary, NC, USA) was used to measure fluorescence in each well at 545-20/600-40 nm wavelength.

Next, to test osmotic stress tolerance of our strains we used two different stressors: polyethylene glycol (PEG) and sodium chloride (NaCl, salt). We repeated the assay described above with different concentrations of Polyethylene Glycol 8000 or NaCl in the R2A medium. Specifically, *Luteibacter* strains were grown at 10% (w/v), 30% (w/v), and 40% (w/v) PEG and 0.3% (w/v), 1.8% (w/v), and 3% (w/v) NaCl.

### Quantitative assay analysis

For every growth condition tested, each biological replicate (strain) was represented by three technical replicates per experimental 96-well plate. Each batch plate growth test was performed in duplicate, resulting in an initial total of six technical replicates for each biological replicate in each growth condition tested. Technical replicates were removed from the dataset if there was clear evidence of contamination in the uninoculated (“blank”) wells associated with specific plate readings. If two or more of the three blank control wells on any experimental plate showed contamination, all readings from that plate were excluded from analysis. For all retained plate readings that passed quality control, relative fluorescence units (RFU) values for each well were corrected by subtracting the average RFU of the uncontaminated blank wells on the same plate. Next, every negative RFU value was assessed individually to determine if there was a technical issue or if there was no cell growth in that well. For strains at each concentration, negative values observed in 2 or more wells were considered biologically meaningful rather than technical errors, changed to zero to indicate no cell growth, and included in the downstream analyses. If a strain only had one negative-value well, this suggested a technical issue and the reading was removed from the dataset. After quality filtering based on blank and negative-value wells, the number of technical replicates retained per biological replicate ranged from two to six, with the majority of strains retaining all six technical replicates for all growth assays (Supplemental Table S2). If a strain had ≤1 technical replicate reading for a growth condition, it was removed from the analysis of that growth condition (Supplemental Table S3)

For all retained plate readings that passed quality control, technical replicate RFUs for each strain within each growth condition were then averaged to yield a single value per biological replicate for downstream analysis.

Using R (v4.5.0) (49), we used a linear model to assess differences in bacterial RFUs in standard R2A media among strains isolated from the three distinct sites (WK, TLI, and KNZ). One-way analysis of variance (ANOVA) was performed using stats::anova() (v4.5.0) to test for significant effects of site on sqrt(RFU). Estimated marginal means (EMMs) were calculated for each site using the emmeans package (v1.11.2) (50), and pairwise comparisons were conducted with Tukey’s Honest Significant Difference (HSD) test. *P-*values were false discovery rate (FDR)-corrected using multcomp::cld() (v1.4-28) (51).

For the stress-tolerance assays, RFU values for each biological replicate under stress conditions were divided by their corresponding RFU values under standard medium control conditions (RFU_stressed_/RFU_control_), calculated separately for each PEG (10%, 30%, 40%) and NaCl (0.3%, 1.8%, 3.0%) concentration. This approach enabled direct comparison of each strain’s change in growth in response to increasing PEG and NaCl concentrations. Differences in the resulting RFU ratio values were then assessed using the same statistical methods described above. Outliers were removed to improve the fit of some models, but did not impact the interpretation of the results.

We assessed *Luteibacter* fitness trade-offs by performing two simple linear regressions of RFU under control conditions on the RFU for both PEG 30% and NaCl 1.8% stress treatments, because these were the conditions under which variation among sites was most apparent (see Results, below). The significance of the observed relationships was evaluated using ANOVA on the fitted linear models using stats::anova() (v4.5.0).

All results were visualized using the ggplot2 package (v3.5.2) (52) in R and Adobe Illustrator was used to create composite figures.

### Whole genome sequencing, assembly, and annotation

Bacterial DNA was extracted from the 96 selected using the Qiagen DNeasy PowerLyzer DNA extraction kit (QIAGEN, Hilden, Germany; #12855-50) and quantified using the QuantiFluor dsDNA kit (Promega, Madison, WI, USA; #E2671) (Supplemental Methods). Bacterial DNA samples were submitted to the Genome Sequencing Core at the University of Kansas for Illumina library preparation and 300 bp paired-end sequencing on an Illumina nextSeq 2000 flow cell (Supplemental Methods).

We received demultiplexed reads from the Genome Sequencing Core. Adapter sequences were trimmed from the paired-end reads and quality filtered using fastp v0.23.4 with the parameters -q 15 -u 40 --trim_poly_g (53). De novo assembly of genomes was completed using SPAdes v3.14 with default parameters in the Shovill v1.1.0 pipeline (54) (Supplemental Table S4). Genome assemblies for each strain were annotated using Bakta v1.9.3 with database v5.1 (55, 56) (Supplemental Table. S5). Raw reads and assembled *Luteibacter* whole genomes are available on NCBI SRA and GenBank under BioProject PRJNA1300453.

### Whole-genome comparisons and pangenome construction

Genome similarity was assessed using MinHash distances calculated with MASH (v2.3) (57) and pairwise average nucleotide identity (ANI) computed using fastANI (v1.34) (58) (Supplemental Table S6 and S7). MASH distance matrices were ordinated in R using classical multidimensional scaling, vegan::cmdscale (v2.7-1) and differences among predefined groups were tested using PERMANOVA (vegan::adonis2) (59).

A pangenome was constructed using Roary (v3.11.2) (60) with a minimum percent identity of 95% under two parameterizations: (i) splitting paralogs and (ii) retaining paralogs to preserve gene copy number information (Supplemental Table S8 and S9). Core gene identity was defined as being represented in 99% of the strains. A single-copy core gene alignment generated by Roary was used to infer a maximum-likelihood phylogeny with IQ-TREE (v2.2.2.6) (61), using the best-fit substitution model selected by Bayesian Information Criterion (GTR+F+G4). The resulting tree was visualized and annotated using iTOL v7.6 (48). Gene presence/absence relationships were visualized in R using ggVennDiagram (v1.5.7) (62).

Gene copy number variation was quantified from the Roary output generated without splitting paralogs. Statistical differences in gene copy number across groups were assessed using Kruskal-Wallis tests (stats::kruskal.test), with Benjamini-Hochberg correction (Supplemental Table S10). Mean copy numbers of significant genes were visualized as heatmaps using pheatmap (v1.0.13) (63).

For strain dereplication, a conservative, phenotype-aware strategy was applied to remove near-identical strains. Strains with >99.99% ANI were evaluated, and redundant strains were removed if they exhibited highly similar phenotypes. Strains with divergent phenotypes were retained despite high genomic similarity. Priority was given to strains with phenotype data for both NaCl and PEG assays. This process resulted in the removal of 20 strains, listed in (Supplementary Table 11), resulting in 76 remaining strains.

### Functional annotation and enrichment analyses

KEGG orthologs (KOs) were assigned to Roary gene clusters (split-paralog run; 12,435 clusters) using KOfamScan with KO profiles and databases obtained from KEGG (downloaded in March 2026) (64, 65). A total of 87% of clusters were assigned KOs, with 6,282 classified as high-confidence assignments. KOs were mapped to KEGG pathways, modules, and BRITE hierarchies using mapping files downloaded from KEGG (April 2026) (Supplemental Table S12). KO assignments were filtered based on KOfamScan thresholds, retaining hits exceeding 75% of the KO-specific score cutoff and with e-values ≤ 1e−50. Annotations corresponding to eukaryotic proteins and non-bacteriophage viral genes (e.g., host signaling proteins, animal receptors, and eukaryotic transcription factors) were manually removed prior to downstream analyses.

For enrichment analyses, counts of gene clusters assigned to each functional category (KO and BRITE B) were summarized per genome and normalized by the total number of annotated terms. Dereplicated strains were used in these analyses. Functional enrichment between semi-arid (SVR, HAY) and humid (KNZ, TLI) climate groups were tested using Kruskal–Wallis tests with false discovery rate (FDR) correction. Significant terms were filtered using the following criteria: (i) adjusted p-value ≤ 0.05, (ii) ≥75% prevalence in at least one group, and (iii) for terms present in 100% of genomes, an absolute mean difference between groups of >2 (Supplemental Tables S13 and S14). To focus interpretation on functional differences nested within broader enriched pathways, significantly enriched KOs were further restricted to those assigned to significant BRITE B categories and classified into ecologically relevant functional groups using a custom in-house script.

### Genome-wide association study

Core genome SNPs were identified using Snippy (v4.6.0), with standard settings (66). Reference-based SNP calling was completed: 1. Across all genomes (reference KNZ12-1B), 2. Lineage I (KNZ/TLI) genomes only (reference KNZ12-1B), 3. Lineage II and III SVR genomes only (reference SVR3-8D). Reference genomes were selected based on assembly quality and phenotype availability. Individual variant calls were combined into a core SNP alignment using snippy-core. Genome-wide association analyses were performed using pyseer (v1.4.1) (37) to identify genetic features associated with osmotic stress tolerance phenotypes (log10 relative growth under NaCl 1.8% and PEG 30% treatments). A fixed-effects model was used and genome structure was controlled for using a MASH distance matrix and standard pyseer settings. Both gene presence/absence (Roary output) and SNP-based analyses were conducted on each of the three dereplicated strain subsets.

Significant SNPs were mapped back to genome annotations (Bakta GFF3 files), and associated loci were linked to Roary gene clusters and corresponding KEGG orthologs. Candidate genes were defined as those with likelihood ratio test (LRT) P-values ≤ 0.05 and absolute β > 1. Because the modest sample sizes (≤76 genomes) provided limited power after multiple-testing correction, the GWAS was treated as a candidate-generating analysis and the unadjusted P-values were used to prioritize loci for future validation. Because of the possible lineage confounding effects, if a SNP/gene was only a significant hit in the all genomes GWAS it was not included as a candidate (Supplemental Table S15).

### Prophage detection and analyses

Prophage regions were annotated from *Luteibacter* whole genome nucleotide sequences using PHASTER (PHAge Search Tool Enhanced Release) webserver (Supplemental Table S16) (67). PHASTER categorized prophage regions as intact, questionable, or incomplete according to the number of phage-associated coding DNA sequences (CDSs) identified within each region, the presence of key phage genes, and similarity to reference prophage genomes. For each predicted prophage region, the genomic coordinates, completeness category, and annotated phage-associated genes were recorded.

Putative phage taxonomy was inferred from PHASTER gene annotations and prophage gene content. Because many prophage regions contained genes with similarity to multiple phage taxa, taxonomic assignments were made conservatively. Prophage regions encoding terminase large subunits, portal proteins, capsid proteins, tail proteins, tape-measure proteins, and other characteristic virion assembly genes were classified as tailed double-stranded DNA phages (class *Caudoviricetes*). In contrast, prophage regions containing filamentous phage replication proteins, gene V proteins, and lacking canonical tailed-phage structural modules, were classified as filamentous phage (family *Inoviridae*). Prophage regions lacking sufficient diagnostic genes were retained as unclassified phage-derived regions.

Next, nucleotide sequences corresponding to all predicted prophage regions were extracted from *Luteibacter* genome assemblies. Prophage sequences were clustered using CD-HIT-EST (68) at 90% nucleotide identity to identify groups of closely related prophages and reduce redundancy. Representative sequences from each cluster were aligned using MAFFT (69) and phylogenetic relationships were inferred using IQ-TREE2 (61) with ModelFinder model selection and 1,000 ultrafast bootstrap replicates together with 1,000 SH-aLRT support replicates.

Prophage comparative analyses were completed using R (70). Near-identical genomes were removed prior to comparative analyses, as mentioned above. Prophage completeness categories (intact, questionable, incomplete, or absent) were visualized for each lineage and site combination using proportional stacked bar plots. The total number of prophage-associated genes predicted by PHASTER was used as a proxy for prophage content within each genome. Differences in prophage gene abundance between strains originating from semi-arid and humid environments were evaluated using Kruskal-Wallis tests.

To determine whether prophage content was associated with osmotic stress tolerance, two linear regression analyses were performed using log-transformed relative growth measurements under NaCl and polyethylene glycol (PEG) stress conditions as response variables and the number of prophage-associated genes as the predictor variable. Separate models were evaluated for NaCl and PEG tolerance phenotypes.

The *Luteibacter* single-copy core gene phylogeny and the prophage phylogeny were compared using cophylogenetic analyses implemented in the R package phytools (71). Trees were rotated to optimize visual correspondence between hosts and prophages, and tanglegrams were generated to visualize congruence between host and prophage evolutionary histories.

## RESULTS

### Soil-dwelling *Luteibacter* spp. vary phenotypically across a climate gradient

We collected 549 *Luteibacter* isolates from a previously described maize root-associated bacterial culture collection (38) that originated from soils across a steep precipitation gradient in Kansas, USA, which is a continuum from semi-arid to humid climate zone, including two semi-arid (SVR and HAY) and humid sites (TLI and KNZ) (Fig. 1a). Only one *Luteibacter* strain was isolated from HAY, therefore this strain was grouped with the SVR strains to represent western Kansas (WK). An initial qualitative phenotyping assay was used to putatively select strains that were tolerant or sensitive to sodium chloride (NaCl), and presumably osmotic stress (Supplemental Fig. 1). Additionally, *16S rRNA* gene diversity of the *Luteibacter* isolates was assessed, and showed that WK strains, apart from the single HAY strain, are in a genetically distinct clade from the humid strains from TLI and KNZ (Supplemental Fig. 2). From the 549 strains, 186 were selected for additional phenotyping, which maximized the phenotypic and genetic diversity of strains from each site.

**Fig 1.**
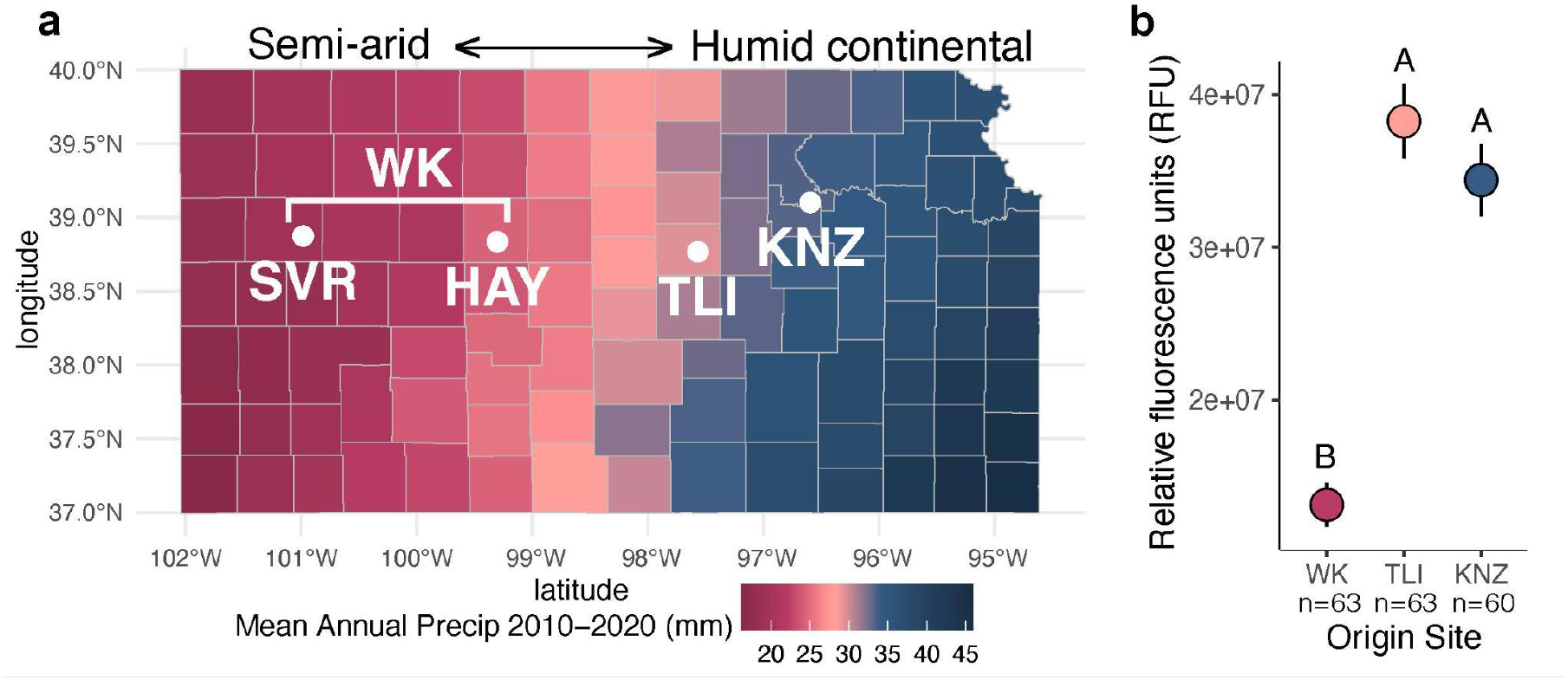
***Luteibacter* strains collected across the Kansas precipitation gradient have different growth potential under controlled laboratory conditions. (a)** Map depicting the mean annual precipitation of Kansas counties for 2010-2020, ranging from a semi-arid to humid continental climate. The four sample sites are SVR, HAY, TLI, and KNZ. SVR and HAY sites were grouped together to represent western Kansas (WK). White points and labels show the locations of the never-irrigated prairie soil sample sites. **(b)** The points represent the estimated marginal means of the relative fluorescence units (RFU), which represents bacterial growth, of the biological replicate strains from each site grown under control conditions. WK strains had significantly less relative growth under controlled conditions compared to TLI and KNZ (ANOVA, F_2,183_=49.117, p<0.0001). Each biological replicate (strain) is the average of 2-6 technical replicates. The error bars represent standard error. Letters represent significance levels (p≤0.05) determined with a Tukey post-hoc test with FDR p-value correction.

Using a high-throughput fluorescence cell viability assay we compared the relative growth of our selected strains in standard R2A liquid medium after 24 hours. Strains from WK grew at <50% the rate of strains isolated from TLI and KNZ (Fig. 1b; ANOVA, F_2,183_=49.12, p<0.001). This indicates that there is a geographical pattern within the phenotypic variation of the *Luteibacter* strains collected.

### Quantitative cell viability assays indicate osmoadaptation fitness trade-offs

Next, we screened the 186 *Luteibacter* strains for osmotic stress tolerance. The cell viability assays were repeated with increasing concentrations of sodium chloride (NaCl) (0.3%, 1.8%, 3.0%) and polyethylene glycol (PEG) (10%, 30%, 40%), and the growth of each strain under stress was normalized to its growth under standard medium conditions to provide the change in growth due to the stress-treatment (relative growth). The WK, TLI, and KNZ strains performed similarly under standard and low-salinity (0.3% NaCl) conditions, with no significant growth reduction (Fig. 2a, NaCl 0.3%; ANOVA, F_2,175_=0.07, p=0.93). However, at moderate salinity (1.8% NaCl), WK strains showed significantly higher relative growth performance than the easternmost KNZ strains, indicating a greater tolerance to salt stress (Fig. 2a, NaCl 1.8%; ANOVA, F_2,182_=3.35, p=0.04). At high salinity (3% NaCl), all strains exhibited markedly reduced relative growth, suggesting a general growth-inhibitory effect at this concentration (Fig. 2a, NaCl 3.0%; ANOVA, F_2,183_=1.98, p=0.14). Under mild osmotic stress (10% PEG), strains had approximately 10-35% relative growth reduction, but there were no significant differences in growth performance among the strains from the three sites (Fig. 2b, PEG 10%; ANOVA, F_2,152_=1.26, p=0.29). However, at 30% PEG, WK strains grew significantly better than TLI and KNZ strains (Fig. 2b, PEG 30%; ANOVA, F_2,148_=3.56, p=0.031). At 40% PEG, the same trend is noticeable, but not significant, suggesting that the strains are approaching a maximum osmotic-stress condition in which all strains perform poorly (Fig. 2b, PEG 40%; ANOVA, F_2,152_=2.14, p=0.12). Together these results confirm that WK strains are better adapted to osmotic stress conditions than KNZ and TLI strains, but are still limited at higher NaCl and PEG concentrations.

**Fig 2.**
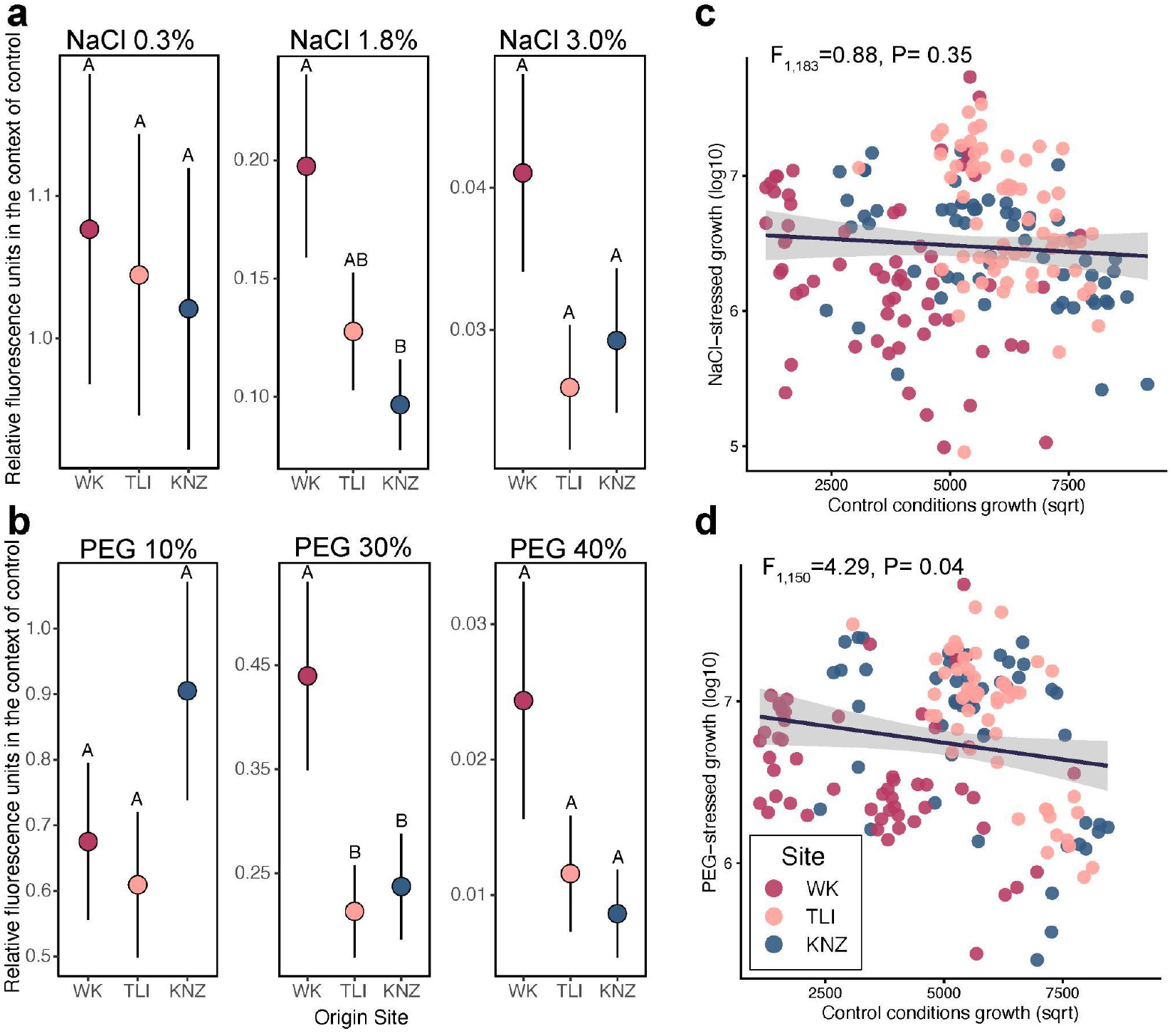
Geographic variation in osmotic stress tolerance is consistent with climate-associated adaptation and reveals a growth–tolerance tradeoff. Relative growth of *Luteibacter* strains under increasing concentrations of NaCl (a) or PEG (b), calculated as RFU under stress divided by RFU under control conditions (RFU_stress_/RFU_control_). Points represent estimated marginal means for strains from each site and error bars indicate standard error. Different letters indicate significant pairwise differences among sites (Tukey post-hoc tests with FDR-adjusted P ≤ 0.05). Sample sizes were: NaCl 0.3%, WK = 55, TLI = 63, KNZ = 60; NaCl 1.8%, WK = 62, TLI = 63, KNZ = 60; NaCl 3.0%, WK = 63, TLI = 63, KNZ = 60; PEG 10%, WK = 54, TLI = 51, KNZ = 50; PEG 30%, WK = 53, TLI = 51, KNZ = 48; and PEG 40%, WK = 54, TLI = 51, KNZ = 50. (c-d) Relationship between growth under control conditions and growth under 1.8% NaCl (C; n = 186) or 30% PEG (D; n = 151). Each point represents one strain, averaged across 2–6 technical replicates, and is colored by site of origin. Black lines show fitted linear regressions with 95% confidence intervals.

We next examined whether growth under standard conditions predicted relative growth under moderate osmotic stress (1.8% NaCl and 30% PEG). For PEG (Fig. 2d, ANOVA, F_1,150_=4.29, p=0.04), but not NaCl (Fig. 2c, ANOVA, F_1,183_=0.88, p=0.35), we observed a significant negative relationship between growth in control conditions and relative growth under stress, consistent with a trade-off between rapid growth and tolerance to PEG-induced osmotic stress.

### Four major *Luteibacter* lineages identified with strong biogeographic genetic patterns

We sequenced the whole genomes of 96 *Luteibacter* strains selected to represent the site, phenotypic, and *16S rRNA* gene diversity observed across the broader isolate collection. Sequencing generated an average of 2,381,658 ± 886,956 300-bp paired-end reads per strain after quality filtering. The assembled genomes averaged 4.3 Mbp across 46 contigs, with an average of 3,721 predicted protein-coding genes per genome (Supplemental Tables S4 and S5). Pangenome analysis using Roary identified 12,438 total gene clusters across the 96 genomes, including only 515 core genes present in ≥95% of strains and 11,923 accessory gene clusters (Supplemental Table S8). This large accessory genome indicates substantial genomic diversity among closely related *Luteibacter* strains.

Core-gene phylogeny and genome-wide distance analyses revealed four major *Luteibacter* lineages, consistent with distinct putative species-level groups based on ANI/MASH distances (Fig. 3a,b, Supplemental Fig. S3, Supplemental Table S6 and S7). These lineages were strongly structured by geography. Lineage I contained strains from the humid sites TLI and KNZ, whereas lineages II, III, and IV were associated with semi-arid sites. Lineage IV contained the single HAY isolate, while lineages II and III both contained SVR strains (Fig. 3a,b). This indicates the genetic divergence was pronounced across sites, but also within semi-arid sites. We identified and removed 20 near-clones (>99.99% ANI and similar assay phenotypes) from the dataset for downstream comparative analyses (Supplemental Table S11).

**Figure 3.**
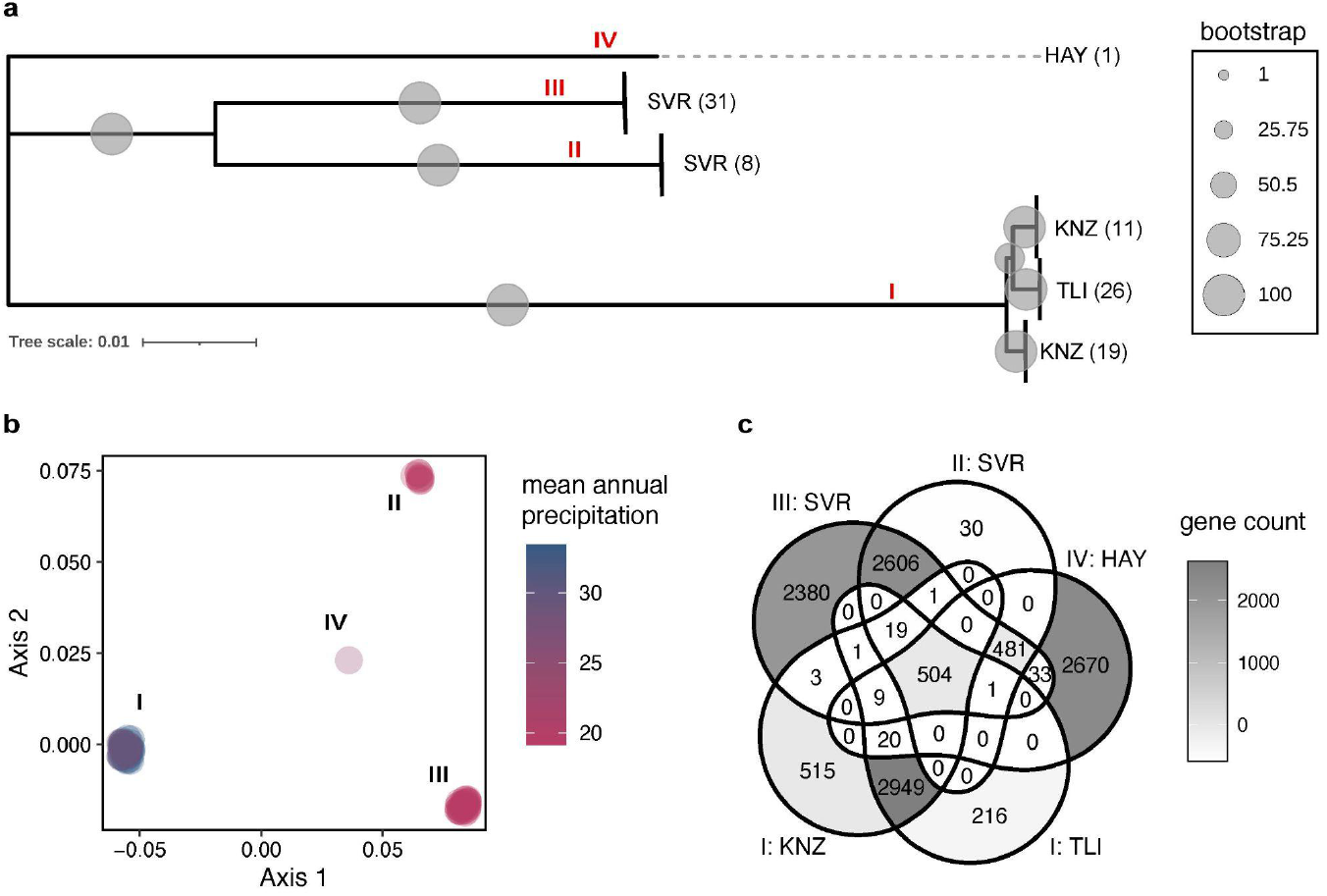
Whole genome analyses reveal strong biogeographic structure among *Luteibacter* lineages. (a) Maximum-likelihood phylogeny constructed from the single-copy core-gene alignment of 96 *Luteibacter* genomes. Four major lineages (I–IV) were identified and were strongly associated with geographic origin: lineage I contained strains from the humid KNZ and TLI sites, lineages II and III contained strains from semi-arid SVR, and lineage IV contained the single HAY strain. Gray circles indicate bootstrap support, with circle size proportional to support value. Branch lengths represent nucleotide substitutions per site. (b) Multidimensional scaling of pairwise MASH genome distances showing separation of the four major lineages. Points represent individual genomes and are colored by mean annual precipitation of the strain’s site of origin. (c) Gene-sharing relationships among the five lineage/site groups (I:TLI, I:KNZ, II:SVR, III:SVR, and IV:HAY) based on the Roary pangenome. Values indicate the number of gene clusters unique to or shared among groups.

Because site and lineage were partially confounded, we next compared gene content across five lineage/site groupings: I:TLI, I:KNZ, II:SVR, III:SVR, and IV:HAY. Gene-sharing patterns further supported strong genomic differentiation among these groups (Fig. 3c). Notably, lineage II:SVR strains had reduced genome sizes relative to the other groups, averaging 117 Kb smaller than lineage III:SVR despite originating from the same site (Fig. 3c, Supplemental Table S4).

Additionally, twenty-one genes had significant copy number variation among lineage/site group (Supplemental Fig. S4). Notably, a gene most closely related to *gp37*, which encodes a *Straboviridae* phage long-tail fiber protein, had an average of 8.1 copies in II:SVR strains, and was absent from all other strains. This lineage-specific expansion suggests that phage-associated genes may contribute to diversification among semi-arid *Luteibacter* lineages.

### Functional enrichment analyses reveal distinct ecological strategies between semi-arid and humid strains

To disentangle broad environmental signatures from lineage-specific effects, we grouped strains by climate of origin rather than sampling site and compared the functional capacities of semi-arid and humid strains. KEGG orthologs (KOs) were assigned to pangenome gene clusters and mapped to hierarchical functional categories, including BRITE B classifications.

The top six BRITE B categories significantly enriched in semi-arid strains were genetic information processing, energy metabolism, xenobiotics biodegradation and metabolism, viral protein families, poorly characterized functions, and glycan biosynthesis and metabolism (Fig. 4a). In contrast, humid strains were enriched in metabolism of amino acids, signaling and cellular processes, signal transduction, carbohydrate metabolism, metabolism, and biosynthesis of other secondary metabolites.

**Figure 4.**
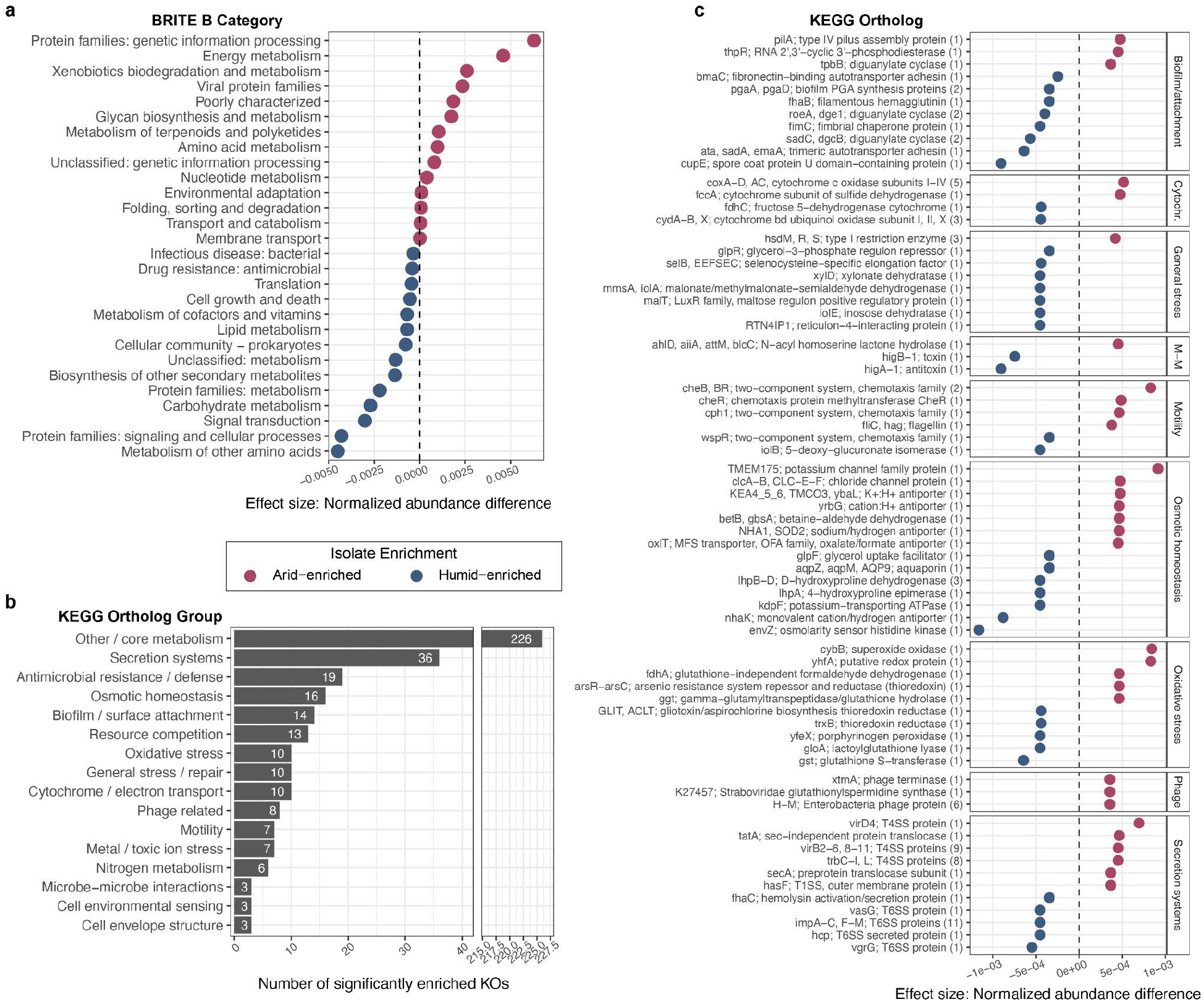
Functional enrichment analyses reveal contrasting genomic strategies between semi-arid and humid continental *Luteibacter* strains. KEGG orthologs (KOs) assigned to pangenome gene clusters were compared between dereplicated strains from semi-arid (SVR and HAY) and humid continental (KNZ and TLI) sites. Significant enrichments were identified using Kruskal–Wallis tests with Benjamini–Hochberg correction (Padj ≤ 0.05) and prevalence filtering. (a) Effect sizes for significantly enriched KEGG BRITE B functional categories. Positive values indicate enrichment in semi-arid strains and negative values indicate enrichment in humid strains. (b) Number of significantly enriched KOs enriched in semi-arid or humid climate regions assigned to manually curated ecological and cellular functional groups. (c) Selected individual KOs illustrating major functional differences between climate groups, including biofilm and attachment, cytochrome/electron transport, general stress, microbe–microbe interactions, motility, osmotic homeostasis, oxidative stress, phage-associated functions, and secretion systems. Points indicate normalized abundance differences, with positive values representing semi-arid enrichment and negative values representing humid enrichment; the dashed line indicates no difference between climate groups. Numbers in parentheses indicate the number of KOs represented by grouped gene annotations.

To further resolve these differences, we manually grouped significantly enriched KOs from the significant BRITE B categories into ecologically relevant functional categories using an in-lab designed script (Fig. 4b). Apart from other/core metabolism-related KOs, the most differentially enriched functional categories were bacterial secretion systems, antimicrobial resistance, and osmotic homeostasis (Fig. 4b).

Several major differences in microbial interaction and colonization strategies emerged between the climate groups. Semi-arid strains were enriched with 18 KOs related to the type IV secretion system (T4SS), whereas humid strains were enriched with 14 KOs associated with the type VI secretion system (T6SS) (Fig. 4c). Humid strains were also enriched in biofilm formation and attachment-related KOs, including *pgaA*, *pgaB*, *fhaB*, and *cupE*, while semi-arid strains were enriched in motility-and chemotaxis-related KOs, including *cheB*, *cheBR*, *cheR*, *cph1*, *fliC*, and *pilA* (Fig. 4c). Additionally, semi-arid strains were enriched with eight phage-related KOs and a N-acyl homoserine lactone hydrolase KO involved in quorum quenching through degradation of quorum-sensing signaling molecules (72).

Major differences were also observed in respiratory and energy metabolism systems. Semi-arid strains were significantly enriched in genes encoding cytochrome c oxidase subunits (*coxA-D* and *coxAC*), whereas humid strains were enriched in genes encoding cytochrome bd ubiquinol oxidase subunits (*cydA*, *cydB*, and *cydX*) (Fig. 4c). These differences suggest divergent respiratory strategies across climate regions.

Semi-arid strains were strongly enriched in functions associated with osmotic homeostasis and ion transport, including Na+/H+ and K+/H+ antiporters, chloride channels, potassium transport proteins, and betaine osmoprotectant biosynthesis pathways (73)(Fig. 4c; Supplemental Table S14). These included genes related to compatible solute production (*betB*/*gbsA*), cation transport (*NHA1*, *yrbG*, *KEA4-6*), and membrane ion flux systems (*clcA-B*, *TMEM175*), collectively suggesting enhanced capacity to maintain intracellular osmotic balance under water-limited conditions.

Semi-arid strains were also enriched in oxidative stress protection and redox-associated pathways, including glutathione metabolism (*ggt*), thioredoxin-linked systems (*arsR-arsC*), superoxide oxidase activity (*cybB*), and formaldehyde detoxification genes (*fdhA*) (Fig. 4c). Additionally, type I restriction-modification systems (*hsdM*, *hsdR*, and *hsdS*) were enriched in semi-arid strains.

In contrast, humid strains were enriched in pathways associated with metabolic versatility and environmental sensing, including carbohydrate utilization, glycerol metabolism, inositol degradation, and osmolarity-responsive regulatory systems (Fig. 4c). Humid strains also encoded osmotic response genes, such as *envZ*, *kdpF*, and aquaporins, although these are more strongly associated with regulatory and transport functions rather than the broader ion-homeostasis systems enriched in semi-arid strains.

Lastly, semi-arid strains were enriched with genes associated with nitrogen metabolism, including *nac*, *ntrY*, *zraR*, *sugE*, and *norR*, whereas humid strains were enriched with genes related to resource competition, particularly iron acquisition systems (*TC.FEV.OM*, *fecR*, *entD*, *hmuT-V*, *ABC.FEV.A*, *fecI*, and *STAR1-2*) (Supplemental Fig. S5). Notably, *TC.FEV.OM* exhibited the largest effect size among all significantly enriched KOs. Together, these results indicate that semi-arid and humid *Luteibacter* lineages differ not only in stress tolerance capacity, but also in broader ecological strategies related to resource acquisition, microbial interactions, and environmental adaptation.

### Genome-wide association study identifies genetic candidates driving *Luteibacter* osmoadaptations

To identify putative genetic determinants of bacterial osmotic stress tolerance, we conducted genome-wide association analyses using both gene presence/absence and SNP-level variation across multiple dereplicated strain sets. Because we have divergent lineages, semi-arid and humid strains could have unique strategies contributing to osmotic tolerance differences within climate regions. Analyses were performed separately within semi-arid strains (SVR only), humid strains (KNZ and TLI), and across all strains (excluding near-clones), for both NaCl 1.8% and PEG 30% relative growth phenotypes (log10 transformed). Additionally, we included MASH distance matrices in the GWAS models to control for population structure.

Relative growth under PEG and NaCl stress was strongly positively correlated across strains (Fig. 5a), indicating that these assays capture a shared axis of osmotic stress tolerance. The strains selected for whole genome sequencing spanned the phenotypic gradient, making them suitability for association analyses.

**Figure 5.**
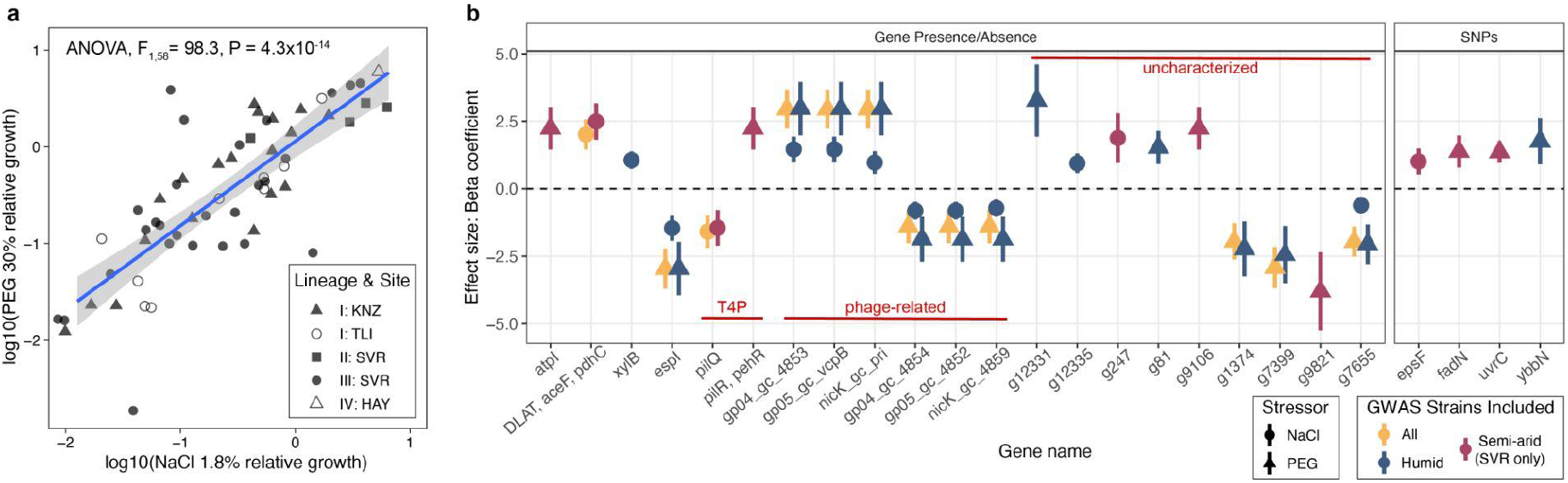
Genome-wide association analyses identify candidate genes associated with variation in Luteibacter osmotic stress tolerance. (a) Relationship between log10-transformed relative growth under 1.8% NaCl and 30% PEG stress for dereplicated strains included in the GWAS. Points represent individual strains and shapes indicate lineage/site group. (b) Effect sizes (beta coefficients ± standard error) for candidate loci identified by gene presence/absence and SNP-based GWAS. Candidates met the thresholds of likelihood ratio test P ≤ 0.05 and |βeta| > 1 and were retained only when supported by a climate-specific GWAS subset. Point shape indicates the stress phenotype tested (NaCl or PEG), and point color indicates the strain subset used in the GWAS: semi-arid (SVR only), humid (KNZ and TLI), or all dereplicated strains. Positive beta coefficients indicate associations with increased.

**Table 1.**
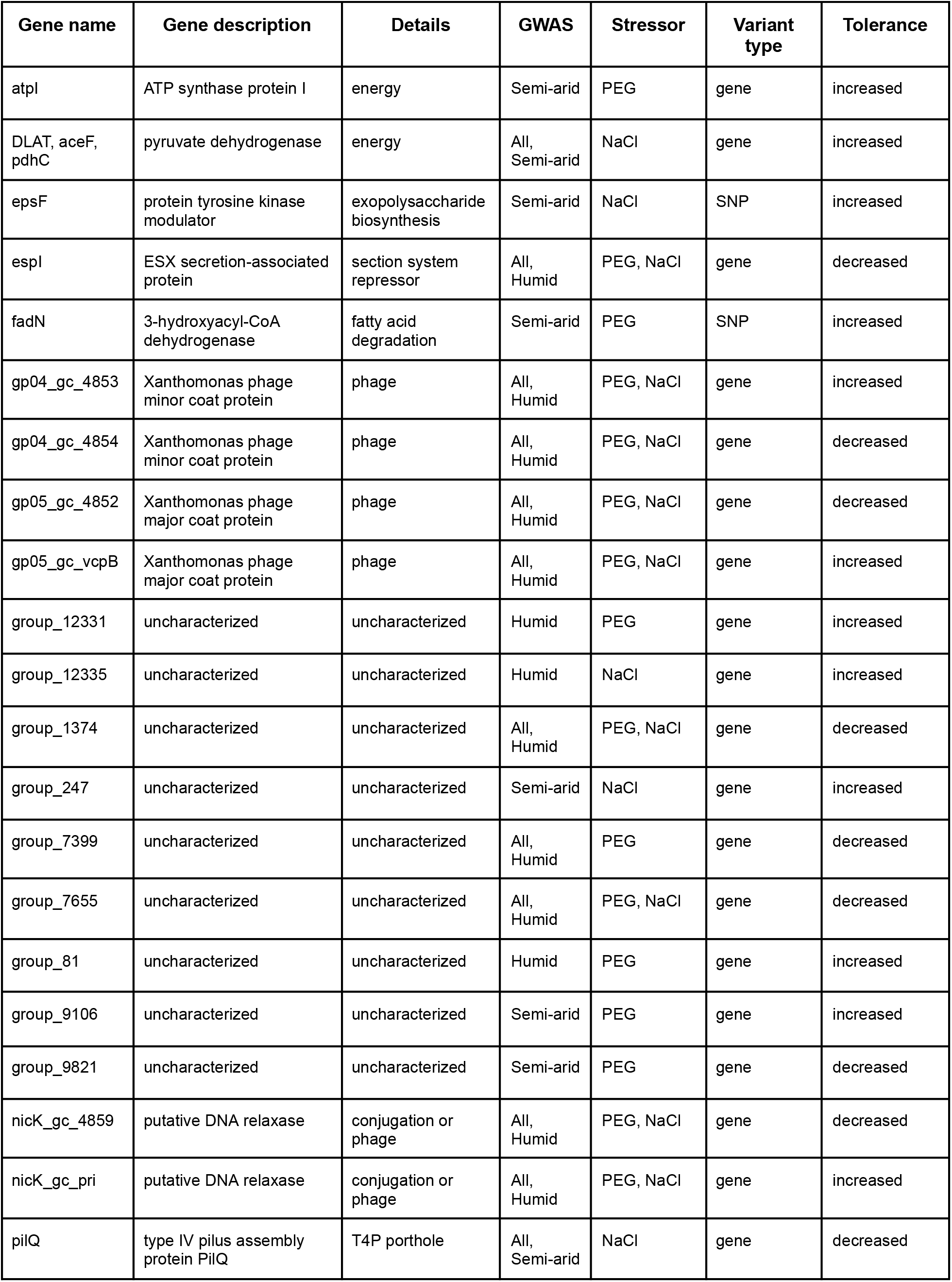
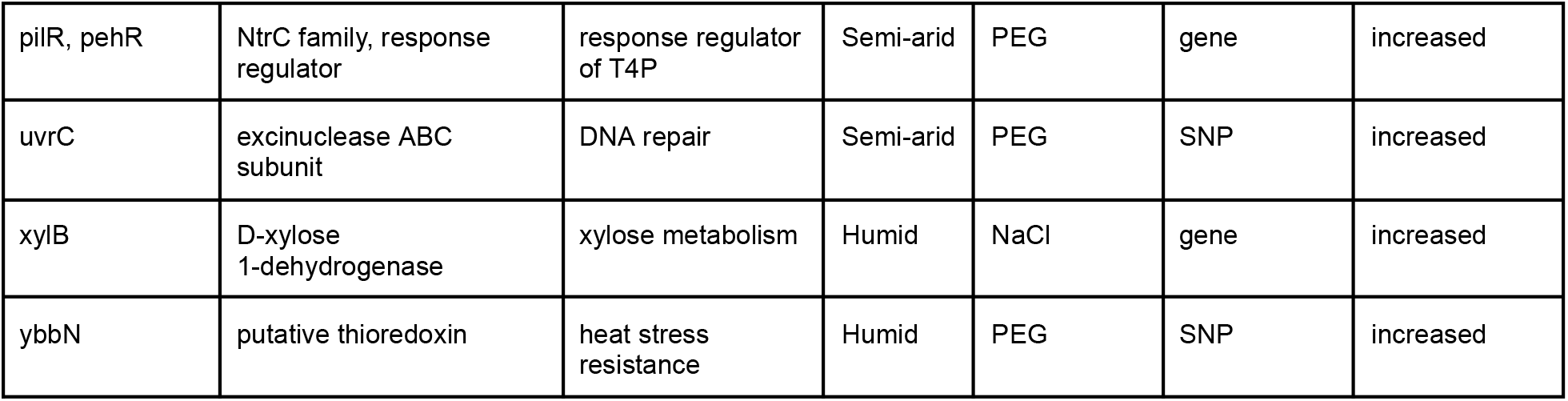
GWAS candidate genes associated with variation in *Luteibacter* osmotic stress tolerance.

| Gene name | Gene description | Details | GWAS | Stressor | Variant type | Tolerance |
| --- | --- | --- | --- | --- | --- | --- |
| atpI | ATP synthase protein I | energy | Semi-arid | PEG | gene | increased |
| DLAT, aceF, pdhC | pyruvate dehydrogenase | energy | All, Semi-arid | NaCl | gene | increased |
| epsF | protein tyrosine kinase modulator | exopolysaccharide biosynthesis | Semi-arid | NaCl | SNP | increased |
| esrI | ESX secretion-associated protein | secretion system repressor | All, Humid | PEG, NaCl | gene | decreased |
| fadN | 3-hydroxyacyl-CoA dehydrogenase | fatty acid degradation | Semi-arid | PEG | SNP | increased |
| gp04_gc_4853 | Xanthomonas phage minor coat protein | phage | All, Humid | PEG, NaCl | gene | increased |
| gp04_gc_4854 | Xanthomonas phage minor coat protein | phage | All, Humid | PEG, NaCl | gene | decreased |
| gp05_gc_4852 | Xanthomonas phage major coat protein | phage | All, Humid | PEG, NaCl | gene | decreased |
| gp05_gc_vcpB | Xanthomonas phage major coat protein | phage | All, Humid | PEG, NaCl | gene | increased |
| group_12331 | uncharacterized | uncharacterized | Humid | PEG | gene | increased |
| group_12335 | uncharacterized | uncharacterized | Humid | NaCl | gene | increased |
| group_1374 | uncharacterized | uncharacterized | All, Humid | PEG, NaCl | gene | decreased |
| group_247 | uncharacterized | uncharacterized | Semi-arid | NaCl | gene | increased |
| group_7399 | uncharacterized | uncharacterized | All, Humid | PEG | gene | decreased |
| group_7655 | uncharacterized | uncharacterized | All, Humid | PEG, NaCl | gene | decreased |
| group_81 | uncharacterized | uncharacterized | Humid | PEG | gene | increased |
| group_9106 | uncharacterized | uncharacterized | Semi-arid | PEG | gene | increased |
| group_9821 | uncharacterized | uncharacterized | Semi-arid | PEG | gene | decreased |
| nicK_gc_4859 | putative DNA relaxase | conjugation or phage | All, Humid | PEG, NaCl | gene | decreased |
| nicK_gc_pri | putative DNA relaxase | conjugation or phage | All, Humid | PEG, NaCl | gene | increased |
| pilQ | type IV pilus assembly protein PilQ | T4P pore | All, Semi-arid | NaCl | gene | decreased |
| pilR, pehR | NtrC family, response regulator | response regulator of T4P | Semi-arid | PEG | gene | increased |
| uvrC | excinuclease ABC subunit | DNA repair | Semi-arid | PEG | SNP | increased |
| xylB | D-xylose 1-dehydrogenase | xylose metabolism | Humid | NaCl | gene | increased |
| ybbN | putative thioredoxin | heat stress resistance | Humid | PEG | SNP | increased |

Candidates had a likelihood ratio test (lrt) p-values ≤0.05 and an absolute beta >1. Using this threshold we identified a set of 18 gene-level candidates and 4 SNP-associated genes (Fig. 5b; Supplemental Table 15). Several candidate genes are directly linked to central metabolism and energy production, including, *atpI*, encoding a component of ATP synthase (74), and the pyruvate dehydrogenase complex (*DLAT*, *aceF*, *pdhC*) (75), which were associated with increased tolerance in semi-arid strains. Additionally, a SNP in *epsF* within semi-arid strains, involved in exopolysaccharide biosynthesis, was associated with increased NaCl tolerance. Similarly, *pilQ* and the *pilR/pehR* regulatory system, involved in type IV pilus (T4P) assembly and regulation, were associated with altered tolerance, implicating surface structures and environmental sensing in stress responses (76).

Genes involved in fatty acid metabolism (*fadN*) and DNA repair (*uvrC*) were also identified in semi-arid strains. In humid-associated strains, genes such as *xylB* (carbohydrate metabolism) and *ybbN* (thioredoxin-related stress response) were linked to tolerance, suggesting alternative strategies for coping with osmotic stress.

Notably, several phage-associated genes, including multiple encoding *Xanthomonas* phage coat proteins gp4 and gp5, and putative DNA relaxases (nicK), were identified in two forms, with one form associated with positive and the other with negative effects on tolerance in humid strains only.

A substantial proportion of candidate loci were annotated as uncharacterized, highlighting the extent of unexplored genetic contributions to bacterial osmoadaptation.

### Predicted phage regions in *Luteibacter* strains are geographically structured

Lastly, we examined patterns of prophage content across the dereplicated *Luteibacter* genomes. Prophage regions were identified using PHASTER and classified as intact, questionable, or incomplete.

Prophage composition varied substantially across *Luteibacter* lineages and geographic origins (Fig. 6a, Supplemental Fig. S6). Most prophage regions identified in lineage I strains from KNZ and all TLI strains were classified as *Caudoviricetes*-like prophages (Supplemental Fig. S6, Supplemental Table S16). In contrast, 16 KNZ lineage I strains contained prophage regions most consistent with filamentous *inovirus*-like phages. Semi-arid SVR strains typically contained two distinct prophage regions: one *Caudoviricetes*-like prophage and a second highly degraded or taxonomically unresolvable phage region. A subset of SVR strains (n=8), corresponding to lineage II:SVR, lacked detectable prophage regions entirely.

**Figure 6.**
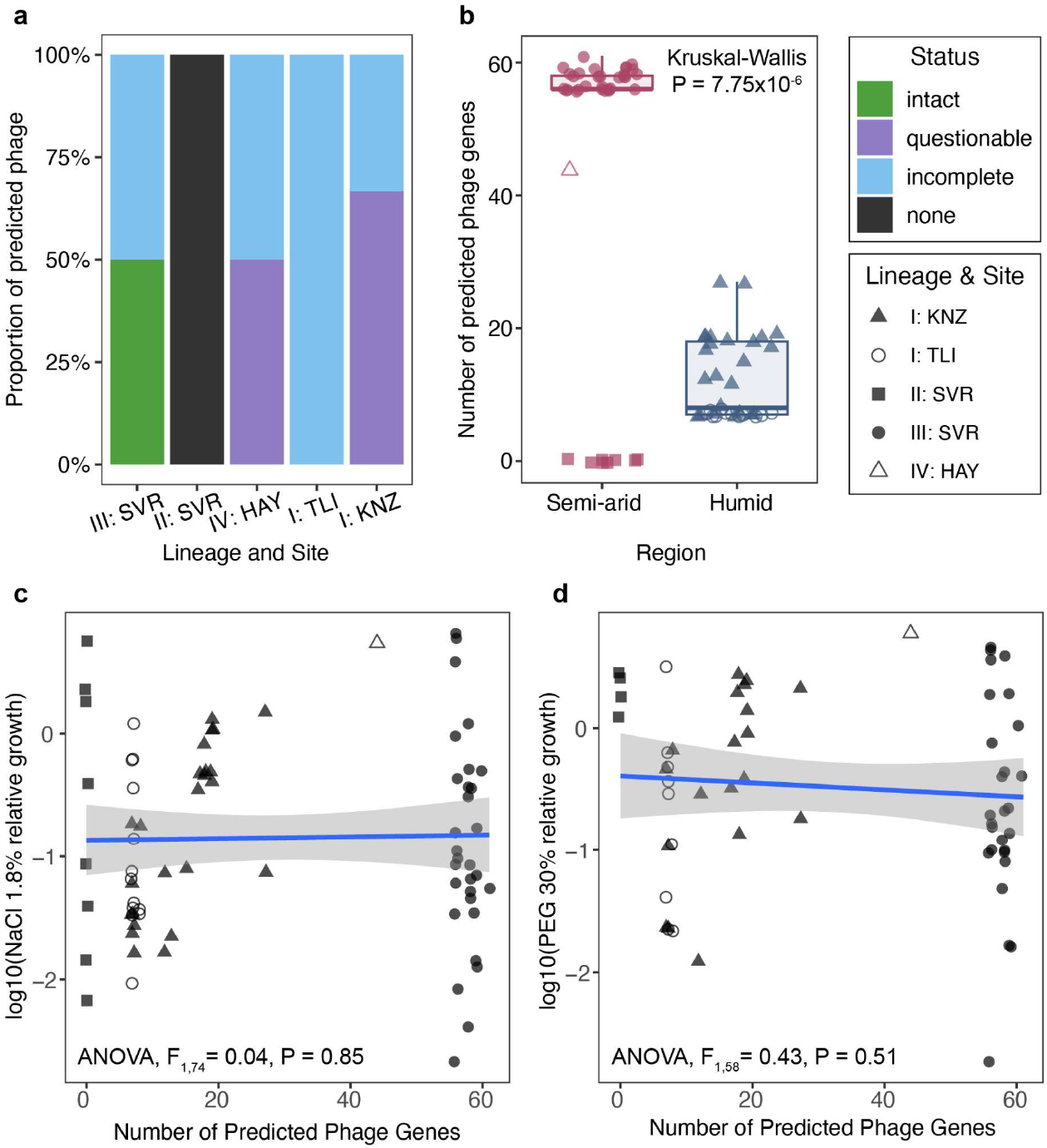
Prophage content is geographically structured and elevated in semi-arid *Luteibacter* strains. (a) Proportion of predicted prophage regions classified by PHASTER as intact, questionable, incomplete, or absent within each lineage/site group. (b) Number of prophage-associated genes predicted per genome in dereplicated strains from semi-arid (SVR and HAY) and humid continental (KNZ and TLI) sites. Points represent individual strains, with shapes indicating lineage/site group; boxes show the distribution within each climate region. (c–d) Relationship between total predicted phage gene content and log10-transformed relative growth under (c) 1.8% NaCl and (d) 30% PEG stress. Lines indicate fitted linear regressions with 95% confidence intervals.

Consistent with these differences in prophage composition, phage-associated gene content varied significantly among lineage and site combinations (Fig. 6a,b). Semi-arid strains, particularly those from SVR, harbored substantially more prophage-associated genes than humid-origin strains, averaging 45.68 phage genes per genome (range: 0–61) compared with 10.89 phage genes per genome (range: 7–28) in humid strains. Notably, the lineage II:SVR strains lacking detectable prophages also exhibited a lineage-specific expansion of a gene most closely related to the *Straboviridae* long-tail fiber protein Gp37, averaging 8.1 copies per genome, whereas this gene was absent from all other strains (Supplemental Fig S4).

Despite these strong biogeographic patterns, total prophage gene content was not significantly correlated with osmotic stress tolerance under either PEG-or NaCl-induced stress (Fig. 6c,d). Thus, while prophage acquisition and retention appear to be strongly structured by evolutionary history and climate, overall prophage burden does not directly explain variation in osmotic stress tolerance.

## DISCUSSION

Our study demonstrates that soil-dwelling *Luteibacter* spp., isolated across a steep precipitation gradient, exhibit substantial phenotypic and genetic diversity, with strong biogeographical structuring. Semi-arid strains tolerated NaCl-and PEG-induced stress better than humid strains. Comparative genomic and GWAS analyses further suggest distinct ecological and genetic strategies across climate regions, with osmotic-stress tolerance associated with variation in energy metabolism, cell-surface biology, DNA repair, and phage-associated genes. Together, these findings indicate that long-term environmental differences shaped not only bacterial stress physiology, but also the broader ecological strategies that influence persistence in dynamic soil environments.

The biogeographical patterns in *Luteibacter* populations are consistent with trends observed in other environmental bacterial populations, like leaf litter *Curtobacterium* (77), soil-dwelling *Streptomyces* (78), and marine *Prochlorococcus* spp. (79). Together, these works signal that environmental bacterial population diversification is the result of gene flow limitations, driven by both isolation by distance and environmental niches. This bacterial genetic diversification translates into phenotypic divergence. For example, Bouskill *et al.* (2025) compared eight phylogenetically paired bacterial isolates from semi-arid and humid tropical soils and found that semi-arid isolates generally maintained greater growth under osmotic stress (80). Similarly, substantial variation in desiccation survival and transcriptional responses among nine *Curtobacterium* strains demonstrated that even closely related bacteria can employ both shared and lineage-specific mechanisms of drought tolerance (81). Our analysis extends these findings, showing that greater osmotic-stress tolerance across semi-arid *Luteibacter* strains coincides with broad genomic differentiation and distinct putative functional strategies. Although precipitation cannot be isolated as the sole driver of lineage divergence, the correspondence among climate of origin, stress phenotype, and genome content is consistent with long-term environmental selection. Moreover, the coexistence of two putative species (ANI 88.6-88.8%) at the semi-arid SVR site suggests that distinct evolutionary trajectories can persist within the same environment. Reduced water connectivity in dry soils may further isolate already heterogeneous soil microhabitats, limiting dispersal and gene flow and potentially helping explain the greater lineage diversification observed at semi-arid compared with humid sites (10, 82).

Predicted functional differences further suggest contrasting ecological strategies between climate groups. Semi-arid strains were enriched in osmotic homeostasis and betaine biosynthesis together with oxidative-stress protection, T4SS, motility, and chemotaxis, suggesting greater investment in maintaining cellular homeostasis and locating favorable microsites under chronic water limitation. Compatible-solute (e.g. betaine) production provides direct protection against osmotic stress, but imposes metabolic costs (83), consistent with the trade-off we observed between rapid growth and PEG-induced stress tolerance. Likewise, motility and chemotaxis may be advantageous in dry, sandy soils where fragmented water films restrict dispersal but transient hydration creates short windows for movement toward favorable microsites (84, 85). Humid strains instead were enriched in T6SS, biofilm and attachment functions, environmental sensing, metabolic versatility, and iron-acquisition systems, consistent with greater investment in resource exploitation and biotic interactions. The T6SS commonly mediates contact-dependent competition and can contribute to nutrient acquisition (86, 87), while enrichment of high oxygen affinity cytochrome bd oxidase, rather than the cytochrome c oxidase functions enriched in semi-arid strains, could favor respiration in low-oxygen microsites associated with wetter soils (88, 89). Thus, climate history may influence the balance among cellular maintenance, mobility, resource acquisition, and microbial competition.

The GWAS results similarly revealed that osmotic stress tolerance has different genetic mechanisms in semi-arid and humid *Luteibacter* populations. Semi-arid candidates broadly implicated energy metabolism, cell-surface properties, and cellular repair. For example, *DLAT*, encoding a component of the pyruvate dehydrogenase complex, was associated with increased tolerance, consistent with experimental links between pyruvate metabolism and bacterial osmotic-stress responses (75). Variation in *epsF*, involved in exopolysaccharide (EPS) synthesis (90), and T4P-related genes (*pilQ* and *pilR*) were also associated with variation in osmotic stress tolerance. EPS can protect cells from dehydration and retain water in dry soils (91), whereas T4P are involved in motility, attachment, environmental sensing, and host interactions (92). *uvrC*, a gene involved in the removal of damaged DNA (93), was also identified as a candidate contributing to osmotic tolerance. Humid-population candidates instead included genes associated with xylose metabolism (94) (*xylB*) and a thioredoxin-like oxidoreductase (95) (*ybbN*). Interestingly, *ybbN* expression is induced by elevated temperatures in other bacteria (96, 97). Notably, no candidate loci were shared between the semi-arid and humid population-specific analyses, consistent with different genetic routes to osmotic tolerance. Given the modest GWAS sample sizes and strong population structure, however, these loci should be considered candidate mechanisms requiring experimental validation rather than definitive causal determinants.

Phage-associated variation adds another dimension to these climate-associated differences. Soil moisture is increasingly recognized as an important driver of virus-host dynamics. For instance, experimental drying can reduce phage activity and increase lysogenic signatures, whereas rewetting seasonally dry soils can rapidly stimulate lytic activity (98, 99). These observations provide an ecological context for the geographic structure of prophage-associated variation in *Luteibacter*, although our data do not indicate that greater prophage burden itself confers osmotic tolerance. Instead, climate-specific associations involving alternative *gp04*, *gp05*, and *nicK* variants raise the possibility that particular phage-derived alleles or linked mobile-elements influence host physiology (100–102).This possibility is especially intriguing in light of the T4P-associated GWAS candidates. T4P frequently serves as phage receptors, and phage proteins can modify T4P assembly, motility, and susceptibility to superinfection (100, 101). Thus, abiotic stress adaptation and phage-mediated selection may intersect at the bacterial cell-surface, but experimental manipulation of T4P-and phage-associated loci will be needed to determine whether these processes are mechanistically connected in *Luteibacter*.

Our findings emphasize that adaptation to climate in natural bacterial populations extends beyond maintaining cellular function during abiotic stress. Persistence in complex soil communities also depends on acquiring resources, navigating heterogeneous environments, and interacting with neighboring microbes, hosts, and phages. Consequently, bacterial survival likely depends on a repertoire of interacting traits rather than a single stress-tolerance mechanism. Understanding how these traits evolve in wild populations will improve predictions of microbial responses to changing climates and may inform the selection or engineering of microorganisms for agricultural, environmental, and biotechnological applications (17, 103).

## DATA AND CODE AVAILABILITY

Raw sequences and assembled *Luteibacter* whole genomes are available on NCBI SRA and GenBank under BioProject PRJNA1300453. Raw data and code related to these analyses are publicly available on GitHub (https://github.com/nginn001/Luteibacter.paper.analyses) and archived on Zenodo (https://doi.org/10.5281/zenodo.22989670).

## ACKNOWLEDGMENTS

The authors would like to thank Drs. Nicole Lukasko and Jason Stajich for helpful feedback and guidance during the preparation of this manuscript. CR was supported by the National Science Foundation REU site “The Stressed Life of Cells” (DBI-2051128 to LD Timmons). This work was supported by the National Science Foundation award IOS-2016351 to MRW and the USDA NIFA award 2022-67013-36672 to MK and MRW. Research reported in this publication was made possible in part by the services of the KU Genome Sequencing Core. This lab is supported by the National Institute of General Medical Sciences (NIGMS) of the National Institutes of Health under award number P30GM145499. This work was also supported by the Kansas INBRE through the award P20GM103418 from the IDeA Program of the National Institute of General Medical Science, as well as a seed grant from the University of Kansas Center for Genomics.

