## Supplemental Figures and Information for "Wild *Luteibacter* populations exhibit climate-associated divergence and distinct genomic strategies for osmotic stress tolerance"

**This PDF file includes:**

Supplementary Figures S1 to S6

Tables S1 to S16

Supplemental Methods

Supplemental References

### SUPPLEMENTAL FIGURES

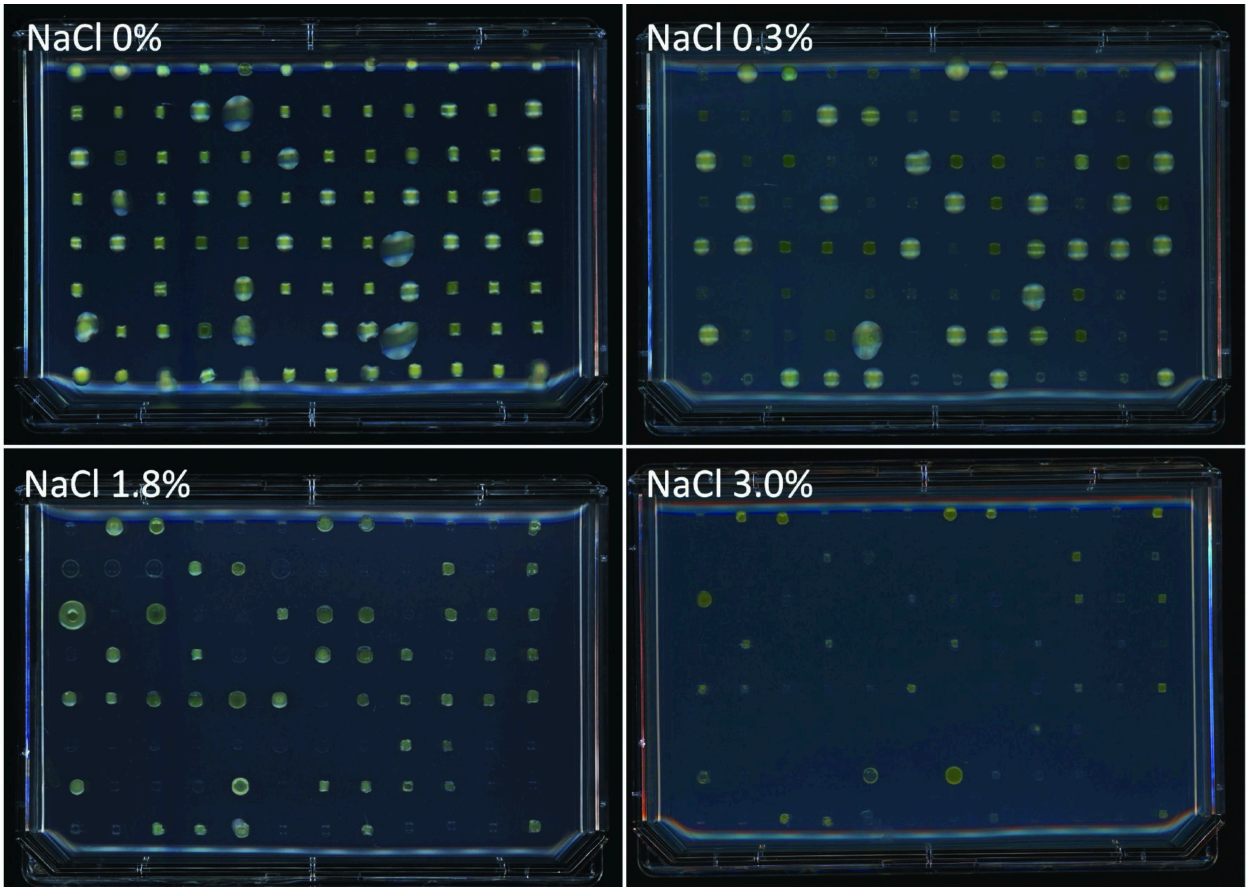


**Supplemental Figure 1. Preliminary qualitative screening of *Luteibacter* osmotic stress tolerance.** Representative growth of *Luteibacter* isolates on R2A agar supplemented with 0%, 0.3%, 1.8%, or 3.0% (w/v) NaCl. Cultures were transferred from randomized 96-well glycerol stocks onto agar plates using a microplate replicator and incubated at 28°C for 48 h. Growth was visually assessed across increasing NaCl concentrations to identify strains with contrasting salt-tolerance phenotypes. Strains exhibiting growth at 1.8–3.0% NaCl were considered relatively tolerant, whereas strains with little or no growth at 0.3% NaCl were considered sensitive. This preliminary screen, together with 16S rRNA gene diversity, was used to select 186 strains for quantitative osmotic-stress phenotyping.

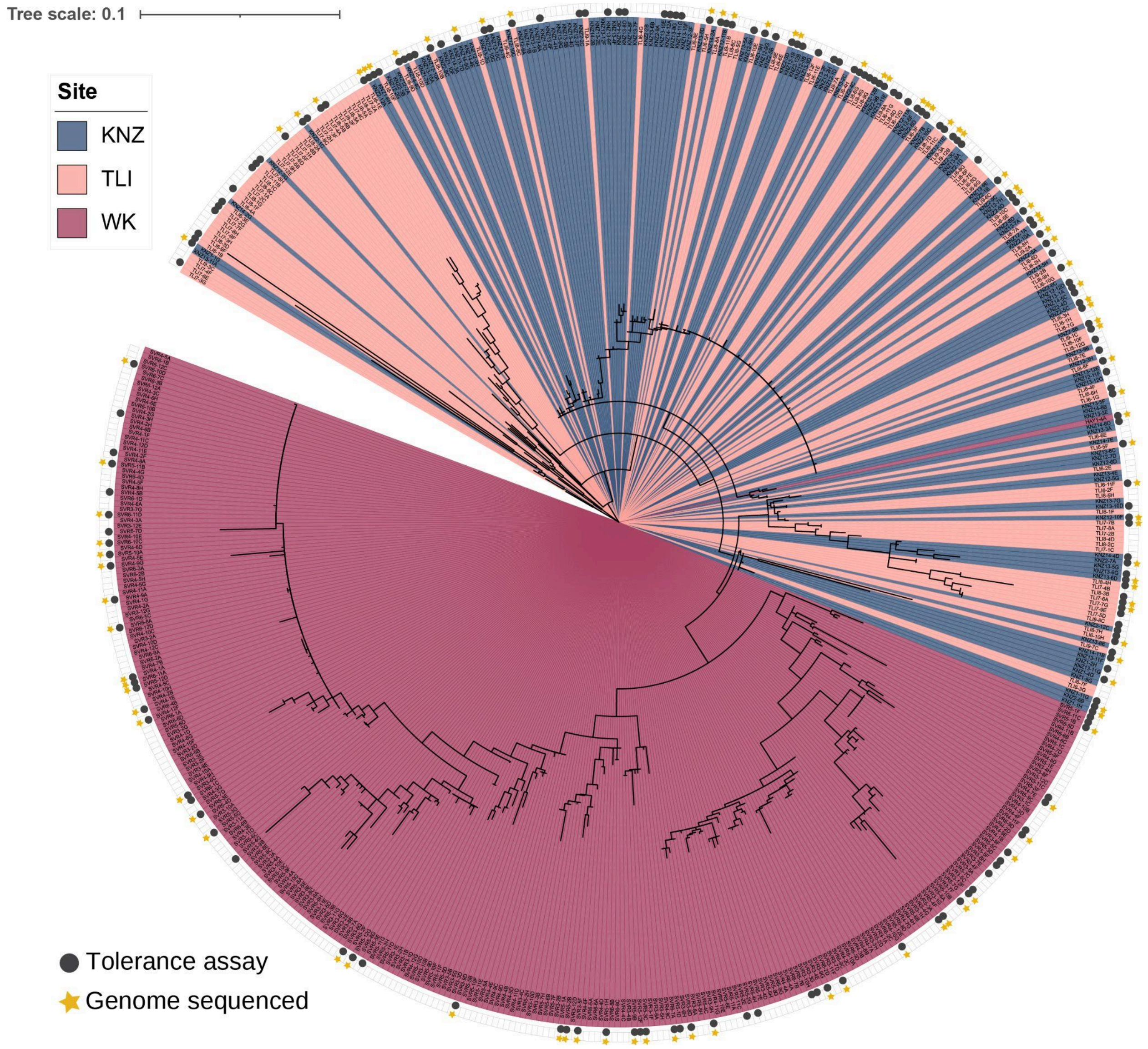


**Supplemental Figure S2. 16S rRNA gene phylogeny of the *Luteibacter* isolate collection.** Maximum-likelihood phylogeny constructed from full-length 16S rRNA gene sequences of 549 *Luteibacter* isolates. Background colors indicate the prairie site of origin: western Kansas (WK; SVR and HAY), The Land Institute (TLI), and Konza Prairie (KNZ). Black circles indicate the 186 isolates selected for quantitative osmotic-stress phenotyping, and yellow stars indicate the 96 isolates selected for whole genome sequencing. Isolates were selected to capture phenotypic and phylogenetic diversity within each site. Branch lengths represent nucleotide substitutions per site.

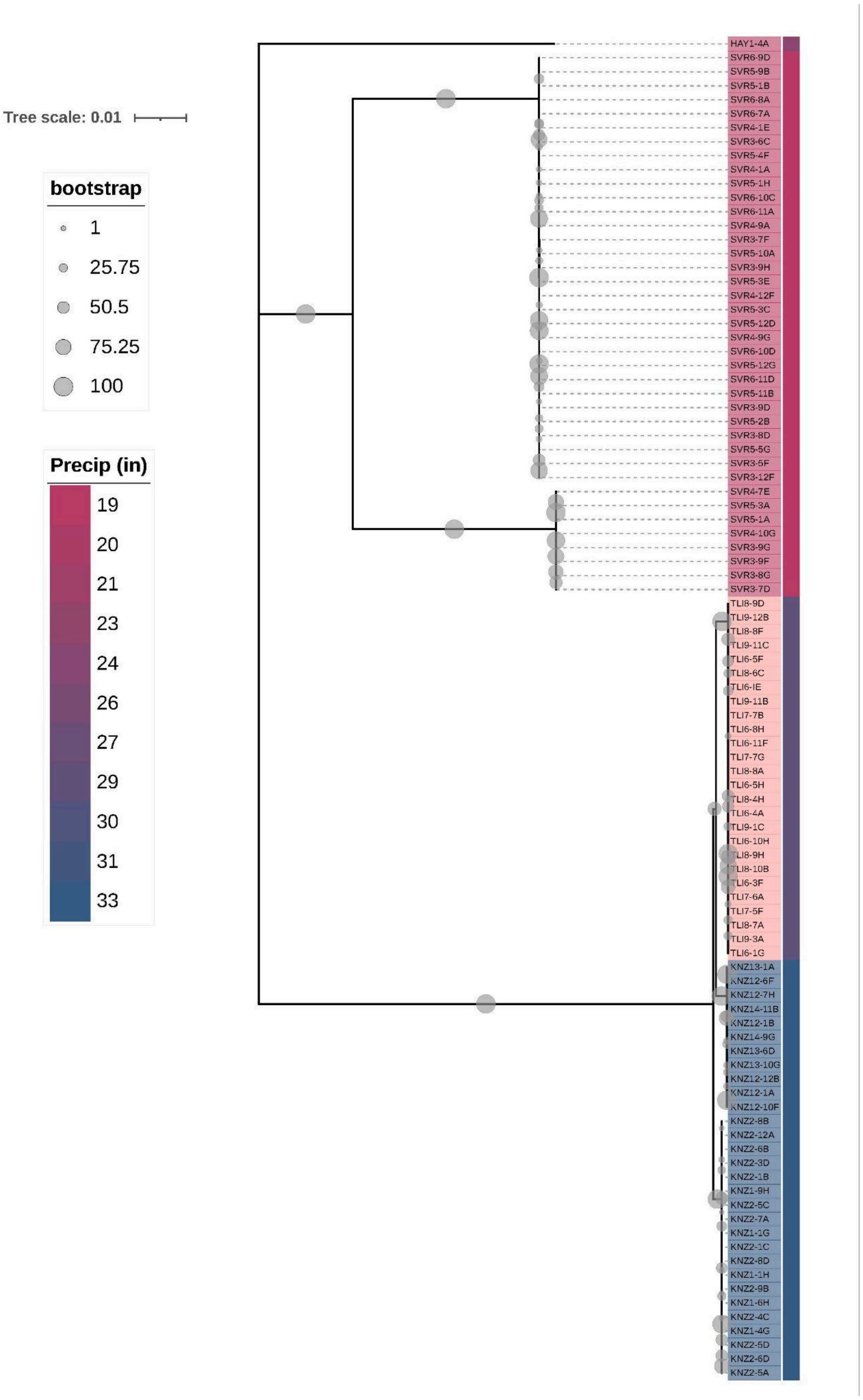


**Supplemental Figure S3. Single-copy core-gene phylogeny of sequenced *Luteibacter* strains.** Maximum-likelihood phylogeny of 96 *Luteibacter* genomes constructed from a single-copy core-gene alignment generated using Roary. Core genes were defined as those present in ≥99% of genomes, and the phylogeny was inferred using IQ-TREE with the best-fit GTR+F+G4 substitution model. Tip labels are colored according to mean annual precipitation (inches) at each strain's site of origin. Gray circles indicate bootstrap support, with circle size proportional to support value, and branch lengths represent nucleotide substitutions per site.

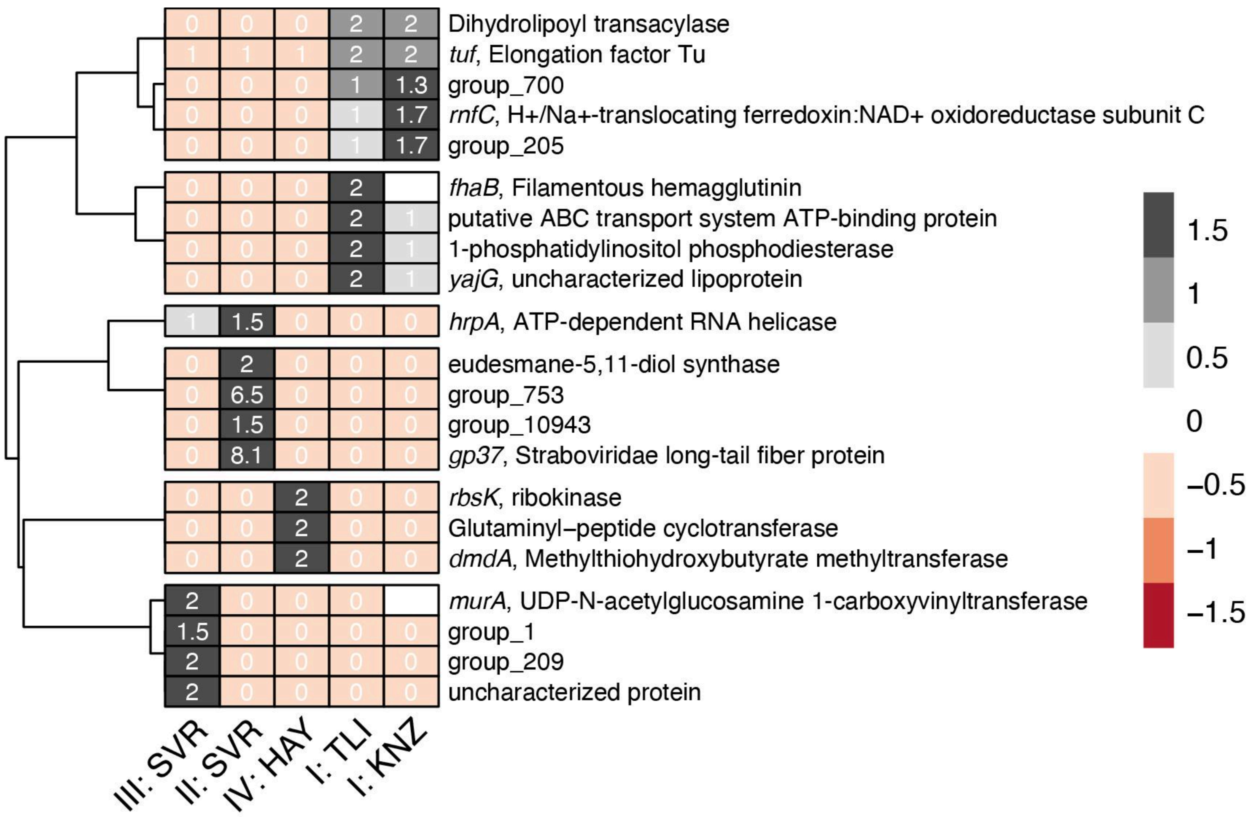


**Supplemental Figure S4. Gene copy number variation among *Luteibacter* lineage/site groups.** Heatmap showing 21 genes with significant differences in copy number among the five lineage/site groups (I:TLI, I:KNZ, II:SVR, III:SVR, and IV:HAY) after removal of near-clonal strains (Kruskal–Wallis test, Benjamini–Hochberg adjusted $P < 0.05$). Colors represent standardized gene copy number (z-score) across groups, whereas values within cells indicate the mean raw gene copy number for each lineage/site group. Gene labels show the top KEGG ortholog annotation and corresponding description; labels beginning with “group” indicate uncharacterized gene clusters.

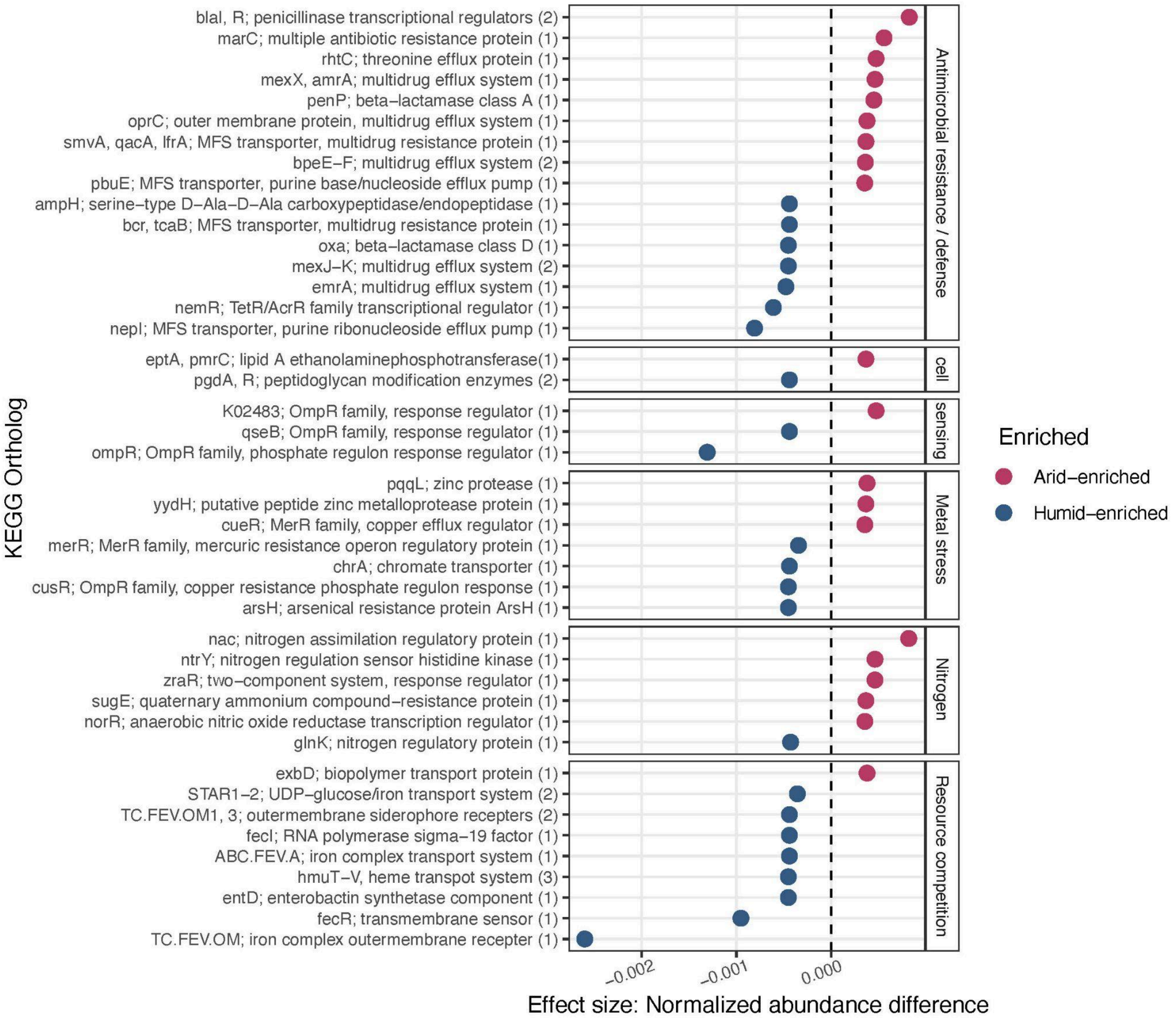


**Supplemental Figure S5. Additional functional differences between semi-arid and humid continental *Luteibacter* strains.** Selected significantly enriched KEGG orthologs (KOs) representing antimicrobial resistance and defense, cell sensing, metal stress, nitrogen-associated functions, and resource competition. Points indicate normalized abundance differences between dereplicated semi-arid (SVR and HAY) and humid continental (KNZ and TLI) strains. Positive values indicate enrichment in semi-arid strains and negative values indicate enrichment in humid strains; the dashed line indicates no difference between climate groups. Numbers in parentheses indicate the number of KOs represented by grouped gene annotations.

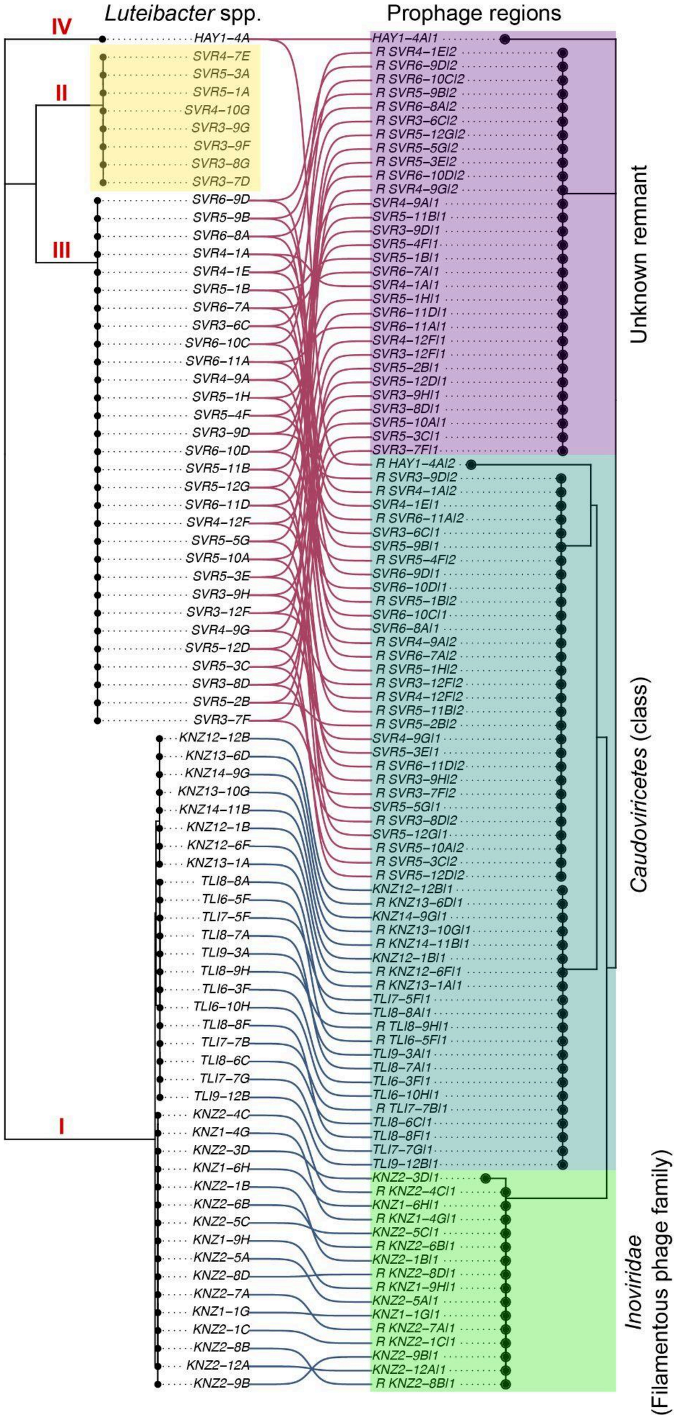


**Supplemental Figure S6. Cophylogenetic comparison of *Luteibacter* hosts and predicted prophage regions.** Tanglegram comparing the *Luteibacter* single-copy core-gene phylogeny (left) with the phylogeny of predicted prophage regions (right) after removal of near-clonal strains. Lines connect each bacterial strain to its corresponding predicted prophage region(s). Host lineages I–IV are indicated on the *Luteibacter* phylogeny, and prophage regions are grouped by putative taxonomy based on PHASTER annotations and diagnostic gene content.

### SUPPLEMENTAL TABLES

**Supplemental Table S1. Kansas precipitation from 2010-2020 by country**.
See attached excel file.

**Supplemental Table S2. Number of technical replicates for each strain in each experiment after quality control.**
See attached excel file.

**Supplemental Table S3. CellTiterBlue assay measurements that passed quality filtering**
See attached excel file.

**Supplemental Table. S4. Genome assembly statistics**
See attached excel file.

**Supplemental Table. S5 Bakta gene annotation information**
See attached excel file.

**Supplemental Table S6. MinHash distances table**
See attached excel file.

**Supplemental Table S7. ANI table**
See attached excel file.

**Supplemental Table S8. Roary pangenome information and counts, split paralogs**
See attached excel file.

**Supplemental Table S9. Roary pangenome information and counts, no split paralogs**
See attached excel file.

**Supplemental Table S10. Gene copy number variation Kruskal–Wallis results**
See attached excel file.

**Supplemental Table S11. Strain metadata table including near-clone information**
See attached excel file.

**Supplemental Tables S12. Full KO/KEGG annotations and count table (split-paralogs)**
See attached excel file.

**Supplemental Tables S13. BRITE B Kruskal–Wallis significant enrichment results**
See attached excel file.

**Supplemental Tables S14. KO with group assignment Kruskal–Wallis significant enrichment results**
See attached excel file.

**Supplemental Table S15. GWAS strong candidates statistics**
See attached excel file.

**Supplemental Table S16. Prophage information and sequences**

See attached excel file.

### SUPPLEMENTAL METHODS

**Prairie soil collection and bacterial isolation collection formation.** Soil was collected from four pristine prairies in Kansas in June 2019 as described in Garrell *et al.* (Garrell et al. 2026), including: The Land Institute (TLI), Konza Wildlife Reserve (KNZ), Hays Prairie (HAY), and Smoky Valley Ranch (SVR). Each of the Kansas sites was split into four subplots. In each subplot, the top ≈10 cm of surface soil and thick plant root masses were removed with a sterilized metal shovel. Then, approximately 1 L of soil was collected and pooled and homogenized in a ziplock bag, for a total of ≈4L of soil collected per site. Soil was held at room temperature for transport back to the laboratory where it was then stored at 4˚C.

Before isolating bacteria from these soils, we conducted one round of plant-association enrichment to increase the relative abundance of organisms that colonize plant roots and, potentially, impact drought tolerance of plant hosts. Maize (*Zea mays* subsp. *mays* var. B73) seeds were sterilized by submerging in 70% ethanol for three minutes, then 5% sodium hypochlorite for two minutes, and finally rinsed in sterile water three times. Sterile seeds were planted in sterile calcined clay (Pro's Choice, #A45216G40, Chicago, IL, USA) in 100 mL cone-tainers or pots. Soil slurries were created by placing 20g of soil in 100ml of 1x PBS with 0.0001% Triton X-100 and shaken at 300 rpms at room temperature for 10 minutes. The soil solutions were filtered through Miracloth into sterile 50mL conical tubes (Midsci #C50B, Valley Park, MO, USA). Tubes were centrifuged for 30 minutes at 3000g. The supernatant was removed and the pellet was resuspended in 20 mL of 1x PBS. The soil slurry was diluted further by adding 5 mL slurry to 500 mL of half-strength MS liquid media. This results in a 0.01 g of soil per mL after dilution. Finally, 25 mL of soil slurry was inoculated into each pot. Pots were then placed in a growth chamber set to a 12-hr day cycle, 27°C/23°C, and ambient humidity for one month.

Roots were collected from one-month old maize plants grown in soil slurries and surface sterilized by being submerged in 95% EtOH and gently shaken for 1 min, then submerged in 3% sodium hypochlorite for 5 minutes, then submerged again in 95% EtOH for 1 minute, and finally rinsed with sterile water three times. Three replicates from the same soil treatment were pooled together and cut into ≤ 1 cm fragments using a flame sterilized razor blade. Between 2-2.5 grams of root fragments for each sample were placed into separate sterile 15 mL conical tubes (MidSci, #C15B, Valley Park, MO, USA). Tubes were then filled with sterile 25% glycerol, incubated at ambient temperature for 20 minutes, and then placed at -20˚C until they were shipped to General Automation Lab Technologies (GALT; San Carlos, CA, USA) for high-throughput microarray isolation using half-strength R2A medium and their Prospector technology, resulting in frozen bacterial liquid cultures preserved in 50% glycerol in a 96-well plate format. Most wells contained a single strain, but some contained mixed cultures. Therefore, each glycerol stock was streaked onto R2A agar medium using a flame sterilized metal loop. Single colonies were selected and subcultured until pure, isolated, individual colonies were obtained.

Bacteria were also isolated from root samples by hand. Briefly, ≈0.5 g of surface sterilized roots were placed into a mesh extraction bag (Agdia, ACC 00930/0100, Elkhart, IN, USA) containing 1 mL of 1x PBS buffer, and pulverized using a hammer. Root slurries were diluted 1:10 with 1x PBS buffer and 100 μl of solution was spread-plate onto R2A medium. Plates were incubated at 28˚C for 5 days. Single colonies were selected and subcultured until pure, isolated, individual colonies were obtained. Pure colonies were then transferred to tryptic soy broth medium in test tubes and incubated for 24 hours at 29˚C shaking at 250 rpm. Liquid cultures of all isolates were preserved in 50% glycerol and stored at -80˚C.

**Bacteria isolate identification and storage.** High-throughput genus-level identification was completed using colony PCR. Briefly, bacterial cells from a single colony were transferred to a PCR tube filled with 20 μl of sterile water using a sterile pipette tip. The cell solution was mixed and incubated at 100˚C for 5 mins using a thermocycler to lyse the cells and release the DNA. The tubes were then placed on ice. The 16S rRNA gene was amplified using in a 25 μL PCR reaction containing 6.5 μL of nanopure sterile water, 2.0 μl of 10μM forward (27F; 5’- AGAGTTTGATCMTGGCTCAG -3’) and reverse (1492R; 5’- ACCTTGTTACGACTT -3’) primers, 12.5 μL of DreamTaq Hot Start PCR Master Mix (2X) (Thermo Fisher Scientific, Waltham, MA, USA; #K9012), and 2.0 μL of DNA template. The thermocycler parameters were 95˚C for 3 mins, 35 cycles of 95°C for 30 s, 50°C for 30 s, and 72°C for 60 s, followed by 72°C for 10 mins. PCR products were run on a 1.0% agarose electrophoresis gel to verify that bacterial amplicons were present. Ten μl of crude PCR product was then mailed to Genewiz (South Plainfield, NJ, USA) for PCR clean-up and Sanger sequencing. Returned sequences were annotated using NCBI Blastn (https://blast.ncbi.nlm.nih.gov/Blast.cgi?PROGRAM=blastn&BLAST_SPEC=GeoBlast&PAGE_TYPE=BlastSearch). From our 1000 bacterial isolates, 549 isolates were classified as part of the genus *Luteibacter*. It is likely that our plant-associated enrichment and isolation methods outlined above enriched for resilient bacterial genera. While our *Luteibacter* strains vary significantly in phenotype, they are all considered fast growers, produce robust biofilms, and are pigmented (yellow)–which offers environmental protection. These traits make them particularly hardy, compared to other soil bacterial genera, and an excellent model for studying environmental stress adaptations.

**Whole genome sequencing**. The 96 strains that were selected were streaked onto fresh R2A plates and incubated at 28°C for 72 hours. A single colony was obtained from each plate and inoculated into a sterile tube filled with 5 mL of R2A liquid medium. Liquid cultures were grown at 30˚C for 48 hours at 250 rpms. For each sample, 1.8 mL of bacterial culture was transferred into a 2 mL microcentrifuge tube. Tubes were centrifuged at 10,000 x g for 1 minute, and the supernatant was discarded. An additional 1 mL of the same culture was added to the pellet, followed by a second centrifugation at 10,000 x g for 1 minute. The supernatant was again removed, and the resulting bacterial pellets were immediately flash-frozen in liquid nitrogen. Pellets were stored at −80°C until extraction. The Qiagen DNeasy PowerLyzer DNA extraction kit (QIAGEN, Hilden, Germany; #12855-50) was used to extract sample DNA according to manufacturer protocol. Extracted DNA was quantified using the Promega dsDNA quantiflour kit (Thermo Fisher Scientific, Waltham, MA, USA; #PRE2671).

Bacterial DNA samples were submitted to the Genome Sequencing Core at the University of Kansas for Illumina NextSeq 2000 Paired End 300 bp library preparation and sequencing. Briefly, gDNA samples were barcoded using unique dual indices, Illumina adapters were added, and the samples were pooled for sequencing.
